# Fate of telomere entanglements is dictated by the timing of anaphase midregion nuclear envelope breakdown

**DOI:** 10.1101/2023.02.24.529917

**Authors:** Rishi Kumar Nageshan, Raquel Ortega, Nevan Krogan, Julia Promisel Cooper

## Abstract

Persisting replication intermediates can confer mitotic catastrophe if left unresolved. Loss of the fission yeast telomere protein Taz1 (ortholog of mammalian TRF1/TRF2) causes telomeric replication fork stalling and in turn, telomere entanglements that stretch between the segregating chromosomes at anaphase. At ≤20°C, these entanglements fail to resolve, resulting in lethality. Rif1, a conserved DNA replication/repair protein, localizes between the segregating chromosomes and specifically hinders the resolution of telomere entanglements without affecting their formation. During anaphase, the spindle and the last segments of segregating chromatin are encased by the nuclear envelope (NE), creating a microdomain termed the anaphase midregion. In the final stages of fission yeast mitosis, this midregion undergoes local NE breakdown. Here we demonstrate that in response to *taz1Δ* telomeric entanglements, Rif1 delays midregion NE breakdown, and this delay disfavors entanglement resolution. Accordingly, gene deletions that hasten midregion NE breakdown phenocopy, and are epistatic with, *rif1+* deletion. Conversely, gene deletions that delay midregion NE breakdown block the *taz1Δ* telomere detanglement afforded by loss of Rif1. Overexpression of Rif1 in a wild type background causes cold-specific NE defects and lethality, which are rescued by treatment with a membrane fluidizing agent. We propose that delayed NE breakdown normally favors the resolution of simple entanglements, arising from incomplete replication, by delaying exposure to the cytoplasm. In contrast, resolution of more complex entanglements involving strand invasion, like entanglements between nonsister *taz1Δ* telomeres, requires more rapid exposure to the cytoplasm. These observations uncover an unexpected coordination between NE remodeling and DNA processing events that can prevent or promote aneuploidy.

## Introduction

A successful mitosis ensures the accurate distribution of the replicated genome to each daughter cell. Each pair of sister chromatids must attach to the mitotic spindle such that the respective sisters are pulled towards opposite poles of the cell. Once chromosome segregation is complete, the spindle disassembles. Mammalian cells undergo an open mitosis in which the nuclear envelope (NE) breaks down completely at mitotic onset, allowing access of a cytoplasmic spindle to the chromosomes^1^. The fission yeast *Schizosaccharomyces pombe* was traditionally considered to undergo a closed mitosis in which the spindle forms and disassembles inside the nucleus. However, two separate instances of local NE breakdown are necessary for this mitosis. First, at the onset of mitosis, NE breakdown beneath the centrosome (called the spindle pole body, SPB) is required to enable SPB insertion into the NE, which is in turn required for formation of the intra-nuclear spindle ^2–5^. Second, the NE breaks down locally in the midregion between segregating chromosomes towards the end of mitosis; the resulting exposure of the late-anaphase spindle to the cytoplasm appears to confer spindle disassembly^6^. Recent studies have documented a sequential dismantlement of specialized nuclear pore complexes (NPCs) that are enriched at the anaphase midregion, and demonstrated that this NPC disassembly triggers the local NE breakdown^7^. Failure to remove NPCs leads to defects in spindle disassembly and failed karyokinesis.

Unresolved replication intermediates that are carried forward to mitosis manifest as chromosome entanglements that are stretched across the anaphase midregion between segregating chromosomes^8–11^. Various repair and replication factors are known to bind and act on these entanglements, but how these factors impact anaphase midregion NE breakdown, and how this NE breakdown affects the fate of entanglements, is unknown^8, 12–15^.

Telomeres, the ends of eukaryotic chromosomes, comprise terminal G-rich repetitive DNA sequences bound by sequence-specific DNA binding proteins and associated proteins, collectively called shelterin, which distinguish the natural ends of chromosomes from those ends produced by damage-induced chromosome breakage^16, 17^. Telomeres also solve the end-replication problem by recruiting the specialized reverse transcriptase, telomerase. Among the shelterin proteins, Taz1 (ortholog of mammalian TRF1/2) specifically binds to telomeric double strand (ds) DNA and forms the foundation for shelterin^18, 19^. In the absence of Taz1, telomeres are deprotected, resulting in telomere fusions and thus dicentric chromosomes that cause mitotic catastrophe ^17, 20, 21^. Telomere fusions are products of classical non-homologous end joining (NHEJ), which occurs only in the G1 stage of the cell cycle. Remarkably, actively growing fission yeast cells have a very short G1 stage and experience virtually no NHEJ^20^. This unique feature of the fission yeast cell cycle enables the investigation of additional functions of Taz1, such as its role in semi-conservative telomere replication, without the complication of chromosome end fusion-mediated cell lethality^22, 23^.

The repetitive G-rich nature of telomeric sequences poses a challenge for replication fork (RF) passage that is relieved by Taz1 binding^22^; mammalian TRF1 plays an analogous role in promoting fork passage^24–27^. In the absence of Taz1, the incoming replisome stalls at telomeres, and processing by sumoylated Rqh1 (the fission yeast RecQ helicase) prevents telomeric RF re-start^28^. As the telomeres at all chromosome ends are homologous and likely to sustain stalled RFs when Taz1 is absent, single strand (ss) DNA from one stalled RF can invade another such RF at a different chromosome end, forming nonsister telomere entanglements^8^. In the ensuing mitosis, these entanglements are stretched between the segregating chromosomes across the midregion, and are resolved during anaphase in cells grown at 32°C but not at temperatures ≤20°C^29^. Hence, *taz1Δ* cells are cold sensitive (hereafter referred to as c/s). Unlike telomere fusions, telomere entanglements are independent of NHEJ. Furthermore, telomere entanglements form in the absence of Rad51, suggesting that telomere-to-telomere strand invasion is independent of standard homologous recombination pathways. Some features of telomere entanglements and how they are processed are apparent from live analysis of mitosis, which reveals the ssDNA binding protein complex RPA bound to telomeric entanglements and interspersed with chromatin; these telomere stretches also harbor DNA polymerase α and Rad52, indicating active processing of the entanglements during mitosis^8^.

Surprisingly, the conserved multifunctional replication/repair protein Rif1 is partially responsible for hindering telomere entanglement resolution and thus conferring *taz1Δ* c/s^8, 30^. First identified as a Rap1 interacting factor in budding yeast, Rif1 controls telomere length and silencing in budding and fission yeast^30–32^. However, as fission yeast Rif1 binds telomeres through Taz1, the role of Rif1 in opposing *taz1Δ* telomeric detanglement is independent of its canonical interaction with telomeres^30, 32^. Crucial nontelomeric roles for Rif1 abound, including the regulation of DNA replication timing and DNA repair pathway choice^33–38^. These Rif1 functions are exerted via its interaction with Protein Phosphatase 1 (PP1) family proteins; indeed, Rif1 can be thought of partly as a scaffold that links PP1 to specific nuclear sites at specific cell cycle stages^36, 39–41^. Loss of Rif1 fails to prevent RF stalling at *taz1Δ* telomeres, nor does it affect the processing of stalled RFs to form telomere entanglements. Instead, the role of Rif1 in *tazΔ* c/s is exerted specifically during anaphase, as demonstrated using the separation-of-function *S-rif1+* allele, which retains Rif1’s S-phase functions but loses its mitotic functions and in doing so, rescues *taz1Δ* c/s^8^.

To delineate the mechanism(s) by which Rif1 inhibits *taz1Δ* telomere detanglement, we performed a series of synthetic genetic array screens to find genes whose deletion reverts the suppression of *taz1Δ* c/s by *rif1Δ* or *S-rif1+*, as well as a more complete compendium of *taz1Δ* c/s suppressors. We find that gene deletions that delay anaphase midregion NE breakdown negate the ability of Rif1 loss to rescue *taz1Δ* c/s. Conversely, gene deletions that advance midregion NE breakdown suppress *taz1Δ* c/s. Accordingly, *taz1Δ* telomere entanglements cause a Rif1-dependent delay in midregion NE breakdown, which in turn influences the cell’s ability to resolve entanglements. Rif1 overexpression leads to cold-specific NE expansion and cell death, which is subverted by treating cells with a membrane fluidizing agent. These results pinpoint the nuclear *versus* cytoplasmic milieus as arbitrators of the final correction mechanisms that ensure chromatin clearance between segregating chromosomes and their successful inheritance.

## Results

### Loss of nuclear pore complex components promotes resolution of telomere entanglements

To investigate the mechanisms that underlie *taz1Δ* entanglement resolution, we performed a series of parallel screens in which the *S. pombe* gene deletion library was crossed with *taz1Δ*, *rif1Δ* and *taz1Δrif1Δ* cells to identify gene deletions that rescue *taz1Δ* c/s or reverse the suppression of c/s conferred by *rif1+* deletion. Based on our previous work, we expected several genetic interactions with *taz1Δ*. For instance, we know that *rap1+* deletion confers synthetic growth defects with *taz1Δ* at temperatures ≤20°C^30^ (note that we routinely incubate cells at 19°C to assess c/s, as indicated hereafter), while *ulp2Δ* is a suppressor of *taz1Δ* c/s^28^. Both of these interactions were recapitulated in our screens (Table S1), validating their efficacy.

Multiple components of the nuclear pore complex (NPC) showed clear genetic interaction with *taz1+* deletion (Figure S1A). Deletions of the nucleoporin-encoding genes *nup60+* or *nup132+* yielded strong suppression of *taz1Δ* c/s. Moreover, independent clones generated by deletion of *nup60+* or *nup132+* rescue *taz1Δ* c/s, validating the screening results (Figure 1A). Epistasis analysis in which *taz1Δrif1Δnup60Δ* triple mutants were compared with *taz1Δnup60Δ* and *taz1Δrif1Δ* double mutants revealed similar degrees of c/s rescue in the double and triple mutants (Figure 1B), suggesting that Nup60 and Rif1 act via a common pathway.

**Figure 1:**
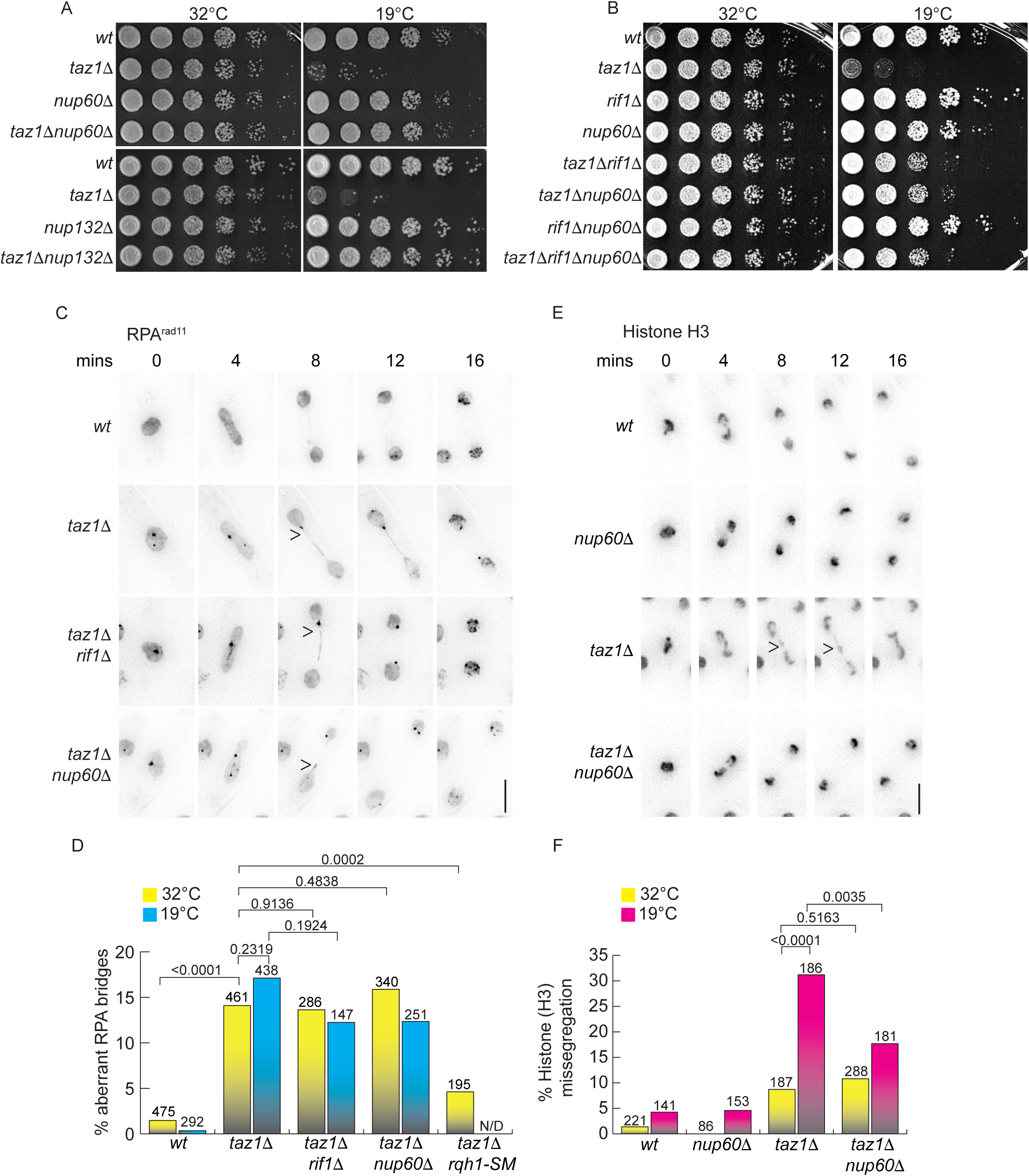
Nuclear pore complex components hinder telomere detanglement: ***A***, 5-fold serial dilutions of log phase (32°C) cells were incubated either at 32°C (2 days) or 19°C (7 days). ***B***, 5-fold serial dilution as in (A). ***C***, Representative images of mitotic cells as visualized by tagging the RPA subunit Rad11 with GFP at its endogenous locus. Aberrant RPA^Rad^^11^ bridges (arrowheads) are seen in *taz1Δ*, *taz1Δrif1Δ* and *taz1Δnup60Δ* cells. An ectopic copy of mCherry-Tubulin (from the *aur1+* locus) was expressed to assess cell cycle stage (not shown). 0’ =metaphase, subsequent increments represent time in anaphase. All time lapse images henceforth are labeled identically. Cells were maintained in log phase at 19°C (3 days) before imaging. Scale bar represents 5μm here and throughout the paper unless otherwise specified. ***D***, Quantitation of aberrant RPA bridges from cells grown either at 32°C (1 day) or 19°C (3 days). n values shown above each bar. N.D =not determined. ***E***, Frames from films of live chromosome segregation visualized via tagged histone H3 (*hht1-RFP*) maintained in log phase at 19°C for 3 days. Arrowheads indicate chromosome missegregation. ***F***, Quantitation of missegregation from films as seen in C grown at 32°C (1 day) or 19°C (3 days). Two tailed Fisher’s exact T test, P values are represented above the brackets in *D* and *F*.

We previously demonstrated that Rif1 affects neither telomeric RF stalling nor stalled RF processing, but rather specifically inhibits the resolution of the consequent telomeric entanglements^8^. A convenient marker of telomere entanglements is the appearance of aberrant bridges that stretch between the separating chromatin masses at anaphase and are bound by the Replication Protein A complex (RPA), a marker of ssDNA. These aberrant RPA bridges often appear punctate and terminate asymmetrically with a concentrated globule of RPA (Figure 1C). These appear in *taz1Δ* cells at both 32°C and 19°C, but are resolved only at 32°C (Figure 1D)^8^. To deepen this analysis of telomeric entanglements, we considered the fact that sumoylated Rqh1 acts on stalled *taz1Δ* telomeric RFs to prevent RF re-start^28^; a sumoylation deficient Rqh1 allele (*rqh1-SM*) enables RF re-start, thereby mitigating *taz1Δ* telomeric entanglement^28^. Hence, while *taz1Δ* cells with proficient *rqh1+* display an increased frequency of aberrant RPA bridges, this frequency is significantly reduced in the *rqh1-SM* background (Figure S1B and 1D), confirming that these RPA bridges are formed from aberrantly processed stalled RFs.

A useful comparison to RPA bridge formation is provided by analysis of bulk chromatin segregation as monitored via a tagged histone (Figure 1E), which reveals chromosome segregation defects downstream of transmission of unresolved entanglements into the next cell cycle^8, 10^. Unlike RPA bridges, which appear in *taz1Δ* cells at 32°C and 19°C, bulk *taz1Δ* chromosome segregation defects are mainly restricted to 19°C (Figure 1F). As we previously demonstrated^8^, *rif1+* deletion has little impact on RPA bridge frequency, confirming that Rif1 affects neither telomeric RF stalling nor stalled RF processing (Figure 1C and D). However, *rif1+* deletion suppresses the chromosome segregation defects seen *via* tagged histones at 19°C, indicating that the aberrant RPA bridges are completely resolved at anaphase^8^ (Figure 1E and F). Thus, Rif1 specifically inhibits the resolution, not the formation, of telomeric entanglements.

If Nup60 and Rif1 act via a common pathway, *nup60Δ*, like *rif1Δ*, would confer *taz1Δ* telomere entanglement resolution without affecting entanglement formation. To investigate this, we compared RPA bridge frequencies in *taz1Δ* cells with and without *rif1+* or *nup60+* at 32°C and 19°C. Consistent with the observed epistatic interactions, loss of *nup60+* suppresses *taz1Δ* c/s without affecting the formation of aberrant RPA bridges (Figure 1C-D). Moreover, Nup60 loss suppresses the appearance of *taz1Δ* chromosome missegregation as observed via tagged histones (Figure 1F), again phenocopying the loss of Rif1. This confirms that the suppression of *taz1Δ* c/s by the loss of Nup60 is not due to averted RPA bridge formation; rather, Nup60 and Rif1 act via a common pathway that regulates entanglement resolution.

### Telomere entanglements cause a Rif1 dependent delay in anaphase midregion NE breakdown

Both Nup60 and Nup132 associate with the nucleoplasmic surface of the NPC and are enriched along the anaphase midregion NE (Figure 2A)^6, 7^, placing them in the cellular domain harboring unresolved telomere entanglements at anaphase. Indeed, the removal of Nup60 and Nup132 is a key step in the localized anaphase midregion NE breakdown^6, 7^. Therefore, suppression of *taz1Δ* c/s by deletion of *nup60+* or *nup132*+ might reflect a role for anaphase NE breakdown in telomere entanglement resolution. To address this possibility, we monitored NPC disassembly and anaphase midregion NE breakdown. Strikingly, NPC disassembly (monitored *via* the loss of Nup60-mCherry signal) in this region is significantly delayed in *taz1Δ* cells at 19°C but not at 32°C (Figure 2B-D). The delay is independent of cell size, as *taz1Δ* cells lacking Chk1, which show normal cell size due to G2/M checkpoint inactivation, show an extent of midregion NE breakdown delay similar to that of *taz1Δ chk1+* cells (data not shown). Crucially, this delay depends on Rif1 (Figure 2B-D).

**Figure 2:**
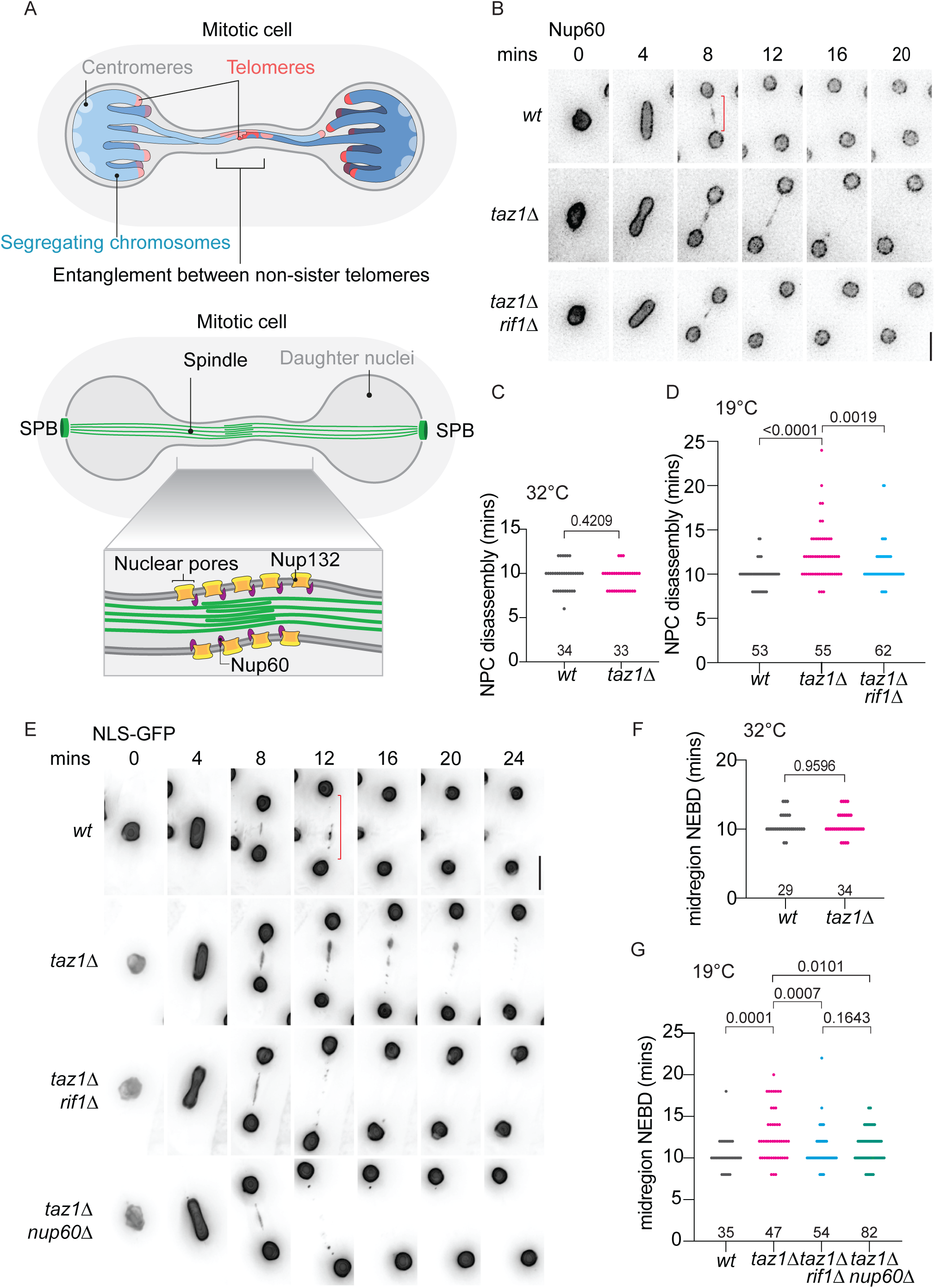
Rif1 delays anaphase midregion NE breakdown in *taz1Δ* cells: ***A***, Schematic of segregating chromosomes harboring telomere entanglements (upper panel), and the anaphase midregion harboring enriched nuclear pore complexes (lower panel). ***B***, Frames from films of mitotic cells expressing Nup60-mCherry from its endogenous locus, maintained in log phase at 19°C. Red bracket represents anaphase midregion. Mitotic progress was monitored via GFP-tubulin (not shown). ***C and D***, The timing of NPC disassembly is plotted relative to anaphase onset from cells maintained either at 32°C (***C***) or 19°C (***D***). ***E***, Frames from films of mitotically dividing cells expressing NLS-GFP-βGAL (a marker of nuclear content whose loss from the nucleus signals loss of NE integrity) maintained in log-phase at 19°C. NLS-GFP-βGAL expression is under control of the *nmt1+* promoter; cells were pre-grown in PMG media without thiamine to induce expression. ***F and G***, Quantitation of midregion NLS-GFP-βGAL signal loss at the indicated time points post-anaphase onset from cells maintained either at 32°C (***F***) or 19°C (***G***). P values derived from Mann-Whitney test is represented above the brackets in C, D, F and G.

To assess NE integrity directly, we utilized NLS-GFP-β-Gal as a marker for nucleoplasmic content; loss of NLS-GFP-β-Gal signal indicates leakage of nuclear contents and thus loss of NE integrity. While *wt* cells show leakage of NLS-GFP-β-Gal approximately 10 minutes after anaphase onset, *taz1Δ* cells show a cold-specific delay (averaging ∼12 mins post-anaphase onset) in midregion NE breakdown (Figure 2E-G). As seen for NPC disassembly, the delay in local NE breakdown is Rif1 dependent (Figure 2G). These observations suggest a model in which Rif1 responds to *taz1Δ* telomere entanglements by conferring a delay in anaphase midregion NE breakdown at 19°C; in turn, this delay stymies resolution of entanglements, leading to Rif1- and Nup60/Nup132-dependent c/s.

### Loss of Mto1 bypasses Rif1’s anaphase function in delaying NE breakdown

To pinpoint the mechanism by which Rif1 impedes telomere detanglement, we probed the screening results for gene deletions that specifically avert the suppression of *taz1Δ* c/s afforded by *rif1Δ*. In parallel with the above-described genetic screens, we performed an analogous series using the separation-of-function allele *S-rif1+*, which abrogates Rif1’s functions in anaphase while preserving its S-phase functions^8^; we selected specifically those gene deletions that confer synthetic sickness in both *taz1Δrif1Δ* and *taz1ΔS-rif1+* backgrounds (Figure S1C and D).

Among gene deletions that specifically restore the c/s phenotype of *taz1Δrif1Δ* and *taz1ΔS-rif1+* cells, *mto1Δ* (encoding <u>m</u>icrotubule <u>o</u>rganizer 1, Mto1) yields a particularly strong synthetic sickness. This genetic interaction was validated by constructing new *mto1+* deletions in independently generated strains of each relevant genotype, confirming that *mto1Δ* specifically reverses the *rif1Δ-* or *S-rif1-*mediated suppression of *taz1Δ* c/s (Figure 3A); *mto1+* deletion has no impact on c/s in the respective single mutant backgrounds. Furthermore, loss of *mto1+* in a *taz1Δrif1Δ* background hinders resolution of telomere entanglements as evinced by increased levels of chromosome missegregation in the triple mutant cells relative to the double mutant (Figure 3B). Indeed, loss of Mto1 reverts histone H3-visualized chromosome missegregation levels to those seen in *taz1Δ* cells at 19°C.

**Figure 3:**
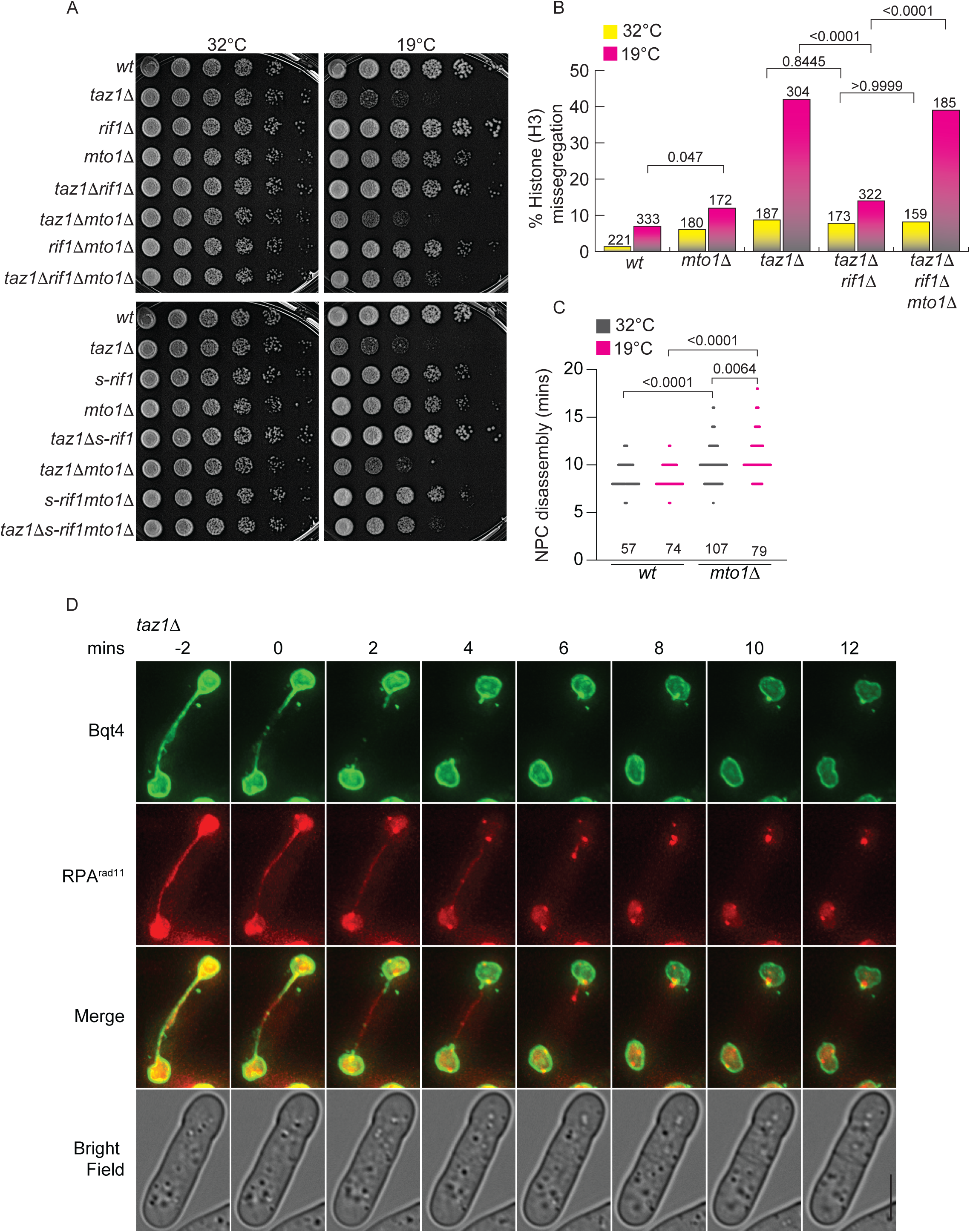
*mto1Δ* reverses suppression of *taz1*Δ c/s by *rif1Δ* by delaying anaphase midregion NE breakdown: ***A***, 5-fold serial dilutions of cells grown to log phase at 32°C were incubated at 32°C (2 days) or 19°C (7 days). ***B***, Quantitation of aberrant histone-H3 missegregation at 32 °C for 1 day and 19°C for 3 days (similar to Figure 1F). For comparison, *wt* and *taz1Δ* 32 °C data is re-plotted from Figure 1F. ***C***, Quantitation of the timing of NPC disassembly time post-anaphase onset from cells maintained either at 32°C for 1 day or 19°C for 3 days. P values from Mann-Whitney test are given above pairs of bars. ***D***, Frames from films of mitotically dividing *taz1Δ* cells maintained in log phase at 32°C. These cells expressed an additional copy of GFP-Bqt4 from its cognate promoter integrated at the *lys1+* locus, while endogenous *bqt4*+ remained unmodified. Rad11-mCherry was expressed from its endogenous locus. Time 0’ marks the onset of anaphase midregion NE breakdown.

### Delayed anaphase midregion NE breakdown mediates the reversion of *rif1Δ* c/s suppression by *mto1Δ*

Mto1 is a component of cytoplasmic microtubule organizing centers and is required for the nucleation of microtubules; it nucleates cytosolic microtubules during interphase, and equatorial and astral microtubules during mitosis^42, 43^. Mto1 also localizes to the SPB during interphase, when the SPB resides on the cytoplasmic surface of the NE; this localization is required for interphase SPB-microtubule interactions^42^. Mto1 mutant alleles that are deficient in forming either equatorial (*mto1-427*) or astral microtubules (*mto1-(1-1065)*) fail to phenocopy *mto1Δ* in a *taz1Δrif1Δ* setting (Figure S1E). Thus, neither disruption of equatorial nor astral microtubules alone is sufficient to confer restoration of *taz1Δrif1Δ* c/s^44^. Moreover, Mto1-(1-1065) fails to localize to the interphase SPB, suggesting that interphase microtubule-SPB interactions are not crucial to *taz1Δ* telomere entanglement resolution.

While Mto1 is dispensable for formation and elongation of mitotic spindles, *mto1Δ* cells show a delay in spindle disassembly at the end of mitosis (see below)^42^. Therefore, we investigated whether anaphase midregion NE breakdown is delayed in the absence of Mto1 by monitoring midregion NPC stability. *mto1Δ* cells show a modest delay in anaphase midregion NPC disassembly at 32°C (Figure 3C) and a profound delay at 19°C. This effect is specific to the *mto1+* null; the *mto1-427* and *mto1(1-1065)* alleles fail to confer a delay in NPC disassembly (Figure S1F). Furthermore, loss of *rif1+* does not impact the delayed NPC disassembly observed in *mto1Δ* cells. Hence, the effect of Mto1 loss in reversing suppression of *taz1Δ* c/s by *rif1+* deletion likely stems from delayed anaphase midregion NE breakdown.

While the *mto1-427* and *mto1(1-1065)* alleles have lost equatorial or astral microtubules, they still possess the post anaphase array^44^, which is lost in *mto1Δ* cells. Indeed, in *wt* cells, we observe networks of cytosolic microtubules crossing the midregion NE bridge at the time of the NE breakdown; these are absent in cells lacking Mto1 (Figure S2A). Hence, Mto1’s role in NE breakdown may involve its nucleation of these NE-disrupting microtubules, which prod and disrupt the midregion NE.

### Midregion NE breakdown is required for telomere entanglement resolution

While *mto1Δ* reverts the suppression of *taz1Δ* c/s conferred by loss of Rif1, *mto1Δ* does not revert the suppression of *taz1Δ* c/s conferred by loss of Nup60 (Figure S2B). Conceivably, loss of Nup60 destabilizes the anaphase midregion NE even in a *mto1Δ* background, thus rescuing the relevant *mto1Δ* phenotype. Indeed, *nup60Δmto1Δ* double mutants display *wt* midregion NE breakdown timing (Figure S2C) while *rif1Δmto1Δ* cells are delayed in midregion NE breakdown. Furthermore, loss of *mto1+* does not change the midregion NE breakdown timing of *taz1Δnup60Δ* cells (Figure S2D). We suggest that the loss of Nup60 guarantees a fragile midregion NE that can break down even in the absence of Mto1-nucleated microtubule prodding.

A conspicuous phenotype of *mto1Δ* cells is spindle persistence. It has been suggested that the lack of cytosolic microtubules in the absence of Mto1 causes an increased concentration of free tubulin, leading to net microtubule polymerization and delaying spindle disassembly^42^. Hence, we considered the possibility that spindle persistence, rather than midregion NE breakdown, is responsible for the inhibition of telomere entanglement resolution. However, loss of Nup60 in a *mto1Δ* background does not rescue spindle persistence even though it rescues *taz1Δ* c/s (Figure S2E). Therefore, c/s rescue stems from the hastened midregion NE breakdown seen in *taz1Δnup60Δmto1Δ* (Figure S2D), not from spindle persistence.

Collectively, these results suggest that cytosolic exposure of entanglements promotes their resolution. Indeed, live analysis of RPA along with a tagged NE protein (GFP-Bqt4) in mitotically dividing *taz1Δ* cells at 32°C reveals that upon midregion NE breakdown (Figure 3D upper panel, see loss of Bqt4 signal), the aberrant RPA bridges are exposed to the cytosol. Only after this exposure are the RPA bridges resolved (Figure 3D RPA panel, also see Video S1-4). The RPA bridge resolution occurs before cytokinesis, allowing the proper chromosome segregation seen in *taz1Δ* cells at permissive temperatures (Figure 3D, lower panel). Thus, telomere entanglement resolution appears to be promoted by cytoplasmic exposure.

### Excess Rif1 causes NE expansion

The anaphase midregion is a well demarcated microdomain that is spatially and temporally restricted^6^. We wondered whether amplification of the effects of Rif1 throughout the nucleus would result in pan-NE defects; therefore, we overexpressed GFP-Rif1 in an otherwise *wt* setting. Remarkably, Rif1 overexpression leads to severe c/s (Figure 4A and S3A). This c/s is likely attributable to Rif1’s actions in mitosis, as *S-rif1+* cells, which overexpress Rif1 by approximately 3-fold in S-phase despite showing 2-fold under-expression in mitosis, grow normally at 19°C. This contrasts with the hypersensitivity of Rif1-overexpressing cells to hydroxyurea treatment, which is shared with the *S-rif1+* allele (Figure S3B) and likely stems from exaggerating the inhibitory function of Rif1 in restraining early origin activation, resulting in genomic under-replication. Our observations are consistent with a recent report that ectopic overexpression of Rif1 from a high copy number plasmid is toxic for *S. pombe* cells^45^.

**Figure 4:**
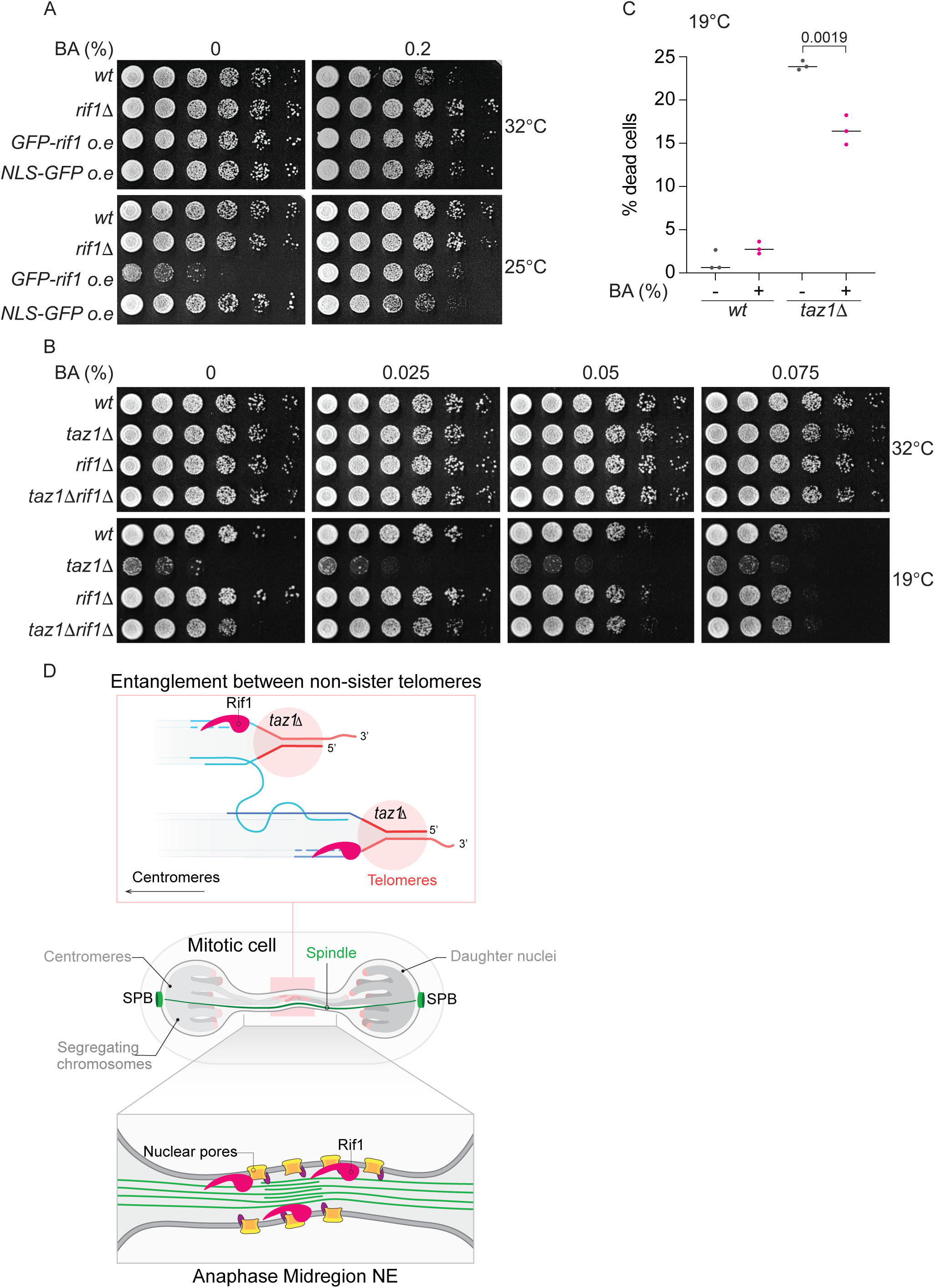
Anaphase Rif1 modulates NE dynamics: ***A***, 5-fold serial dilutions of log phase cells were plated on minimal PMG media (without thiamine to allow induction of *nmt41-gfp-rif1+*, integrated at the endogenous *rif1+* locus) with or without BA and incubated at 32°C (3 days) or 25°C (7 days). ***B***, 5-fold serial dilutions of log phase cells at 32°C were stamped on rich media (YES) containing various concentrations of BA and incubated at 32°C (3 days) or 19°C (10 days). **C**, Quantitation of % of dead cells when grown in media without or with 0.025% BA at 19°C for 3 days. Each data point is result from one experiment scored >1000 cells. Mean value from three independent experiments is represented for each condition. ***D***, Working model for Rif1’s control of anaphase midregion NE breakdown. Rif1 bound telomere entanglements (top) are shown stretched across the anaphase midregion (middle panel). The lowest panel shows Rif1 coordinating the presence of telomere entanglements, which harbor ss/ds DNA junctions that can bind Rif1, with modifications to NE components including Nup60.

To investigate the effects of Rif1 overexpression on the NE, we assessed interphase nuclear morphology via tagged inner-NE and NPC components (Man1-tomato and Nup60-mCherry, respectively). In *wt* cells grown at 19°C, the interphase nucleus is spherical, with Man1 and Nup60 signals evenly spread around the NE (Figure S3C). In contrast, Rif1 overexpression induces NE expansion, with folds of NE appearing and an uneven distribution of Man1 and Nup60 (Figure S3C). This NE expansion is specific to Rif1-overexpressing cells grown at 19°C; no such irregularities are seen at 32°C (Figure S3C). We also observe increased spindle persistence in Rif1-overexpressing cells grown at 19°C (Figure S3D).

The NE expansion seen upon Rif1 overexpression demonstrates that Rif1 alters NE properties in a cold-specific manner. To explore this idea further, we treated Rif1-overexpressing cells with the membrane fluidizing agent benzylalcohol (BA). Fission yeast cells are sensitive to BA at ≤20°C and this is further exacerbated on the minimal medium needed for overexpression of Rif1; to circumvent this lethality, we grew the cells at moderately cold temperature (25°C) where Rif1 overexpression is also near-lethal (Figure 4A). Strikingly, BA treatment rescues the severe c/s induced by Rif1 overexpression (Figure 4A). Thus, Rif1 appears to counteract membrane fluidity, which is required for midregion NE breakdown and in turn, telomere entanglement resolution.

### Membrane fluidizing agent suppresses *taz1*Δ c/s

As BA treatment abates the c/s stemming from Rif1 overexpression, we queried whether it could also rescue *taz1*Δ c/s by opposing the effects of Rif1 on NE rigidity. *Wt* cells lose viability progressively with increasing BA concentration at 19°C, while no such viability reduction is seen with increasing BA concentration in *taz1Δ* cells (Figure 4B); in contrast, BA has no effect on growth at 32°C of either *wt* or *taz1Δ* cells. The near-lethality of BA at ≤20°C made it impossible to confidently interpret dilution assays at this temperature. However, we were able to maintain *wt* and *taz1*Δ cells in log phase growth in the presence or absence of low concentrations of BA at 19°C for three days, after which we checked the percentage of dead cells using Erythrosin B, a dye that stains dead cells. While the percentage of dead *wt* cells is unaffected by BA in liquid culture, the appearance of dead *taz1Δ* cells is partially suppressed by BA (Figure 4C). Likewise, *taz1Δ* cells that are continuously propagated in low concentrations of BA at 19°C show improved viability relative to the same cells without BA (Figure S3E). This suppression is incomplete, likely due to the overall impact of BA on all cellular membranes; in addition, Rif1-controlled pathways likely counteract the effects of BA. Consistent with this notion, BA addition potentiates suppression of *taz1Δ* c/s by *rif1Δ* (Figure 4B), enhancing suppression to the extent of recapitulating *wt* levels of growth. These data establish a key connection between membrane dynamics and the temperature specificity of *taz1Δ* telomere detanglement defects. We propose that NE fluidity at 32°C is optimal for anaphase midregion NE breakdown irrespective of Rif1 status, thereby enabling *taz1*Δ entanglement resolution. However, increased membrane rigidity at 19°C, which is augmented by Rif1’s presence, causes a delay in midregion NE breakdown, which in turn impedes telomere entanglement resolution.

## Discussion

The compartmentalization of the eukaryotic genome within the nucleus prevents exposure of chromosomes to the cytosol through most of the cell cycle. Nonetheless, the limited or complete breakdown of the NE at specific cell cycle stages violates this compartmentalization. Here we find that the resulting exposure of chromatin to the cytoplasm has profound implications for resolving entanglements that occur due to telomeric replication impediments. We find that in response to the presence of *taz1Δ* telomeric entanglements, Rif1 mediates a delay in midregion NE breakdown, therefore prolonging the period during which DNA processing pathways operate without cytoplasmic exposure. Surprisingly, this prolongation hinders the resolution of telomere entanglements. Our data show that Rif1 counteracts NE breakdown by altering the behavior of NE components, including subunits of the NPC (Figure 4D). The effects of Rif1 on nuclear membrane biology shed light on the reasons for the cold-specificity of the lethality caused by Taz1 loss, tying this lethality to the effects of temperature on the biophysical properties of the NE.

### Numerous genetic interactions point to NE breakdown

The convergence of numerous genetic interactions between NE breakdown-associated factors and *taz1+* deletion outlines an intimate relationship between anaphase midregion NE breakdown and telomere detanglement. The loss of NPC components whose disappearance is known to trigger localized anaphase midregion NE breakdown, Nup60 or Nup132, phenocopies and is epistatic with the loss of Rif1 in suppressing *taz1Δ* c/s. Accordingly, *taz1+* deletion leads to a cold-specific, Rif1- and Nup60/Nup132-dependent delay in anaphase midregion NE breakdown.

Our parallel screen for genes whose deletion reverses *rif1Δ*-mediated suppression of *taz1Δ* c/s also converged upon anaphase midregion NE breakdown by revealing that Mto1 is required for suppression by *rif1Δ*. Cells lacking Mto1 both show abnormal spindle persistence and recapitulate the delayed anaphase midregion NE breakdown seen in *rif1+* cells. However, as deletion of *nup60+* in the *mto1Δ* background reverses the NE breakdown delay without reversing spindle persistence, we can separate the two processes and pinpoint the former in controlling *taz1Δ* c/s. Intriguingly, these observations indicate that spindle disassembly is not automatically triggered by cytosolic exposure if free tubulin concentrations are elevated. We speculate that in addition to controlling cellular concentrations of free tubulin, the cytoplasmic microtubules nucleated by Mto1 stimulate midregion NE breakdown by ‘poking’ the midregion NE (Figure S2A). Thus, spindle persistence and quashed resolution of telomere entanglements appear to be two independent phenomena caused by delayed anaphase NE breakdown.

### Confounded replication re-start pathways at *taz1Δ* telomeres lead to perilous anaphase

Interactions between stalled RFs and NPCs in earlier stages of the cell cycle have been implicated in RF restart. In fission yeast, stalled RFs accumulate SUMO-modified proteins, which confer Rad51-dependent relocation to NPCs where processing by the SUMO protease Ulp1 allows fork restart^46^. Likewise, localization to NPCs of stalled telomeric RFs in human cells, or of stalled RFs at CAG repeat regions in budding yeast, during S-phase promotes RF re-start^47, 48^. However, whether such processes are relevant to stalled *taz1Δ* telomeric forks is unclear. Unlike RFs that re-start upon association with NPCs, Rad51 is dispensable for the processing of stalled *taz1Δ* telomeric RFs^29^. Moreover, while telomeres are constitutively positioned at the NE in a Taz1-independent manner, compromised localization of telomeres to the NE has no impact on *taz1Δ* c/s (^28, 49^ and data not shown), suggesting that NPC-driven RF re-start is inactive at *taz1Δ* telomeres. Also of note are our previous observations that sumoylated Rqh1 promotes the aberrant processing of stalled telomeric RFs that prohibits RF re-start, suggesting that in the absence of Taz1, local sumoylation is detrimental rather than beneficial to the completion of chromosomal replication. Conceivably, such RF re-start pathways are specifically detrimental by promoting strand invasion from one stalled telomeric RF into a stalled RF on a non-sister telomere (Figure 4D, upper panel).

### Crucial connections between membrane dynamics and DNA processing events

The involvement of anaphase midregion NE breakdown in *taz1Δ* c/s, the suppression of *taz1Δ* c/s by a membrane fluidizing agent, and the cold-specific NE distortions and NE rigidity-associated lethality triggered by Rif1 overexpression, all reveal a link between the cold-specificity of *taz1Δ* lethality and NE dynamics. The Rif1-mediated delay in NE breakdown in response to telomere entanglements is itself cold-specific. Hence, *taz1Δ* c/s provides a rare window into the profound effects of delayed cytoplasmic exposure on DNA processing events, as the cold-specific alteration of the biophysical properties of the NE reveals the intersection between NE dynamics and DNA entanglement resolution at anaphase. We propose that this interplay is a conserved feature of eukaryotic mitosis. Indeed, depending on specific NE compositions and NPC modification states across different mammalian cell types and growth conditions, Rif1 may regulate mitotic NE dynamics at temperatures greater than 20°C. Moreover, temperatures of ≤20°C are commonly found in the natural habitat of fission yeast and indeed may represent physiological conditions.

Recruitment of Rif1 to the anaphase midregion could be via NE and/or DNA interactions. The N-terminus of Rif1 binds ss-dsDNA junctions^50^, a likely mode of recruitment of Rif1 to *taz1Δ* telomeric entanglements (Figure 4D, upper panel); Rif1’s G-quartet binding capacity may also contribute^35, 51^. However, Rif1 may also associate directly with the NE as suggested from previous studies. In budding yeast, palmitoylation of Rif1 confers NE targeting^52, 53^, which in turn promotes the ability of Rif1 to bind DSBs and favor NHEJ over end-resection^54^. Both human and fission yeast Rif1 associate with Triton-X-100- and DNase-insoluble fractions through biochemical purification protocols^35, 51, 55^ and Rif1 colocalizes with sites of DNA replication at the nuclear periphery in mid-S-phase mammalian cells^33, 34, 38^. Intriguingly, anaphase midregion localization of Rif1 is also widely conserved, being first observed in human cells^56^. Moreover, midbody-bound Rif1 has been shown to regulate the timing of abscission, the final step of cytokinesis, in an anaphase DNA bridge-independent manner^57^. Hence, both NE and DNA association likely concentrate Rif1 at the anaphase midregion, where it regulates NE dynamics, at least in part by regulating PP1-mediated dephosphorylation of NPC components and/or NE phospholipids^8^ (Figure 4D, lower panel). Indeed, a phosphoproteomic analysis of *wt*, *rif1Δ* and *S-rif1+* cells grown at 32°C versus 19°C identified Nup60 as a Rif1-targeted phosphatase substrate. Moreover, the extractability of Nup60 is affected profoundly by Rif1’s presence in *taz1Δ* cells specifically at ≤20°C (Figure S4 and its corresponding supplementary information), suggesting the possibility that Rif1 controls the phosphorylation status of Nup60 and its membrane association.

Human Rif1 was shown to bind non-telomeric ultrafine anaphase bridges and promote their resolution^15^. This positive role for Rif1 in ultrafine anaphase bridge resolution might initially appear paradoxical given our observations that fission yeast Rif1 opposes the resolution of telomeric entanglements. However, we surmise that the Rif1-mediated delay of NE breakdown, as well as any additional roles played by Rif1 in entanglement resolution, has evolved to promote resolution of sister entanglements, the most common type of chromosome entanglement arising from incomplete replication and the likely scenario in cells displaying non-telomeric ultrafine anaphase bridges. Only in rarer scenarios do non-sister entanglements arise, including the *taz1Δ* scenario in which multiple, clustered, telomeres at different chromosome ends will harbor stalled RFs, providing multiple substrates for non-sister strand invasions and the formation of nonsister entanglements, whose resolution is opposed by Rif1^8^.

The crucial question raised by our work is how persistence of the anaphase midregion NE in *rif1+* settings blocks resolution of *taz1Δ* telomeric entanglements. NE breakdown may expose entanglements to cytosolic factors that promote resolution. Alternatively, exposure to the cytoplasm may change or halt the activities of nuclear DNA processing factors that control telomere detanglement. A prime candidate for such regulation is topoisomerase 2 (Top2), which plays an important role in *taz1Δ* telomere entanglement resolution^29, 58^; current experiments aim to address the idea that Top2 activity changes upon exposure to the cytoplasm.

### Conserved mechanisms of DNA processing across a continuum of extents of NE breakdown

While a continuum of the extent of NE breakdown exists across eukaryotes, the closing stages of mitosis are universally followed by the formation of daughter nuclei that should each be encapsulated by a new membrane. In human cells, which undergo complete mitotic NE breakdown, telomere fusions that elicit dicentric chromosomes lead to the formation of NE bridges between daughter cells that persist into the next cell cycle; in the subsequent interphase, bridge rupture leads to untimely cytoplasmic exposure with profound consequences for chromosome segregation, including chromothripsis and kataegis. Moreover, as cells harboring telomere fusions traverse mitosis, the presence of telomeric DNA and RNA in the cytoplasm leads to autophagy and activation of innate immune signaling, respectively^59–64^. Our work demonstrates that different types of telomeric associations trigger different resolution processes. We suspect that mammalian telomere entanglements analogous to the non-sister entanglements we study herein form upon telomeric replication stress. Indeed, we find anaphase RPA-coated bridges upon TRF1 conditional knockout in mouse embryonic fibroblasts, and observe mRif1 localizing to these bridges (data not shown). Conceivably, the presence of mRif1 on those bridges influences whether they become encapsulated in new NE structures and whether such NE structures harbor NPC components^65^. It will also be crucial to determine whether the DNA processing events that govern DNA entanglement resolution are similarly regulated by cytosolic exposure across species.

## Acknowledgements

We thank our lab members, all of whom have contributed vital ideas and advice, Lakshmi Sreekumar for help in double blind experiments (Figure S4C), Michael Lichten (NCI), Dermot Cooper (University of Cambridge), Eros Lazzerini Denchi and his lab (NCI) for crucial discussions, and Jiewei Xu (UCSF) for help in performing synthetic genetic array screens. We thank Ken Sawin (Wellcome Center, Edinburgh) for sharing *mto1+* separation of function mutants. We particularly thank Maria Diaz de la Loza for help with illustrations. This work was supported by the National Cancer Institute and the University of Colorado School of Medicine.

## Author contributions

RKN performed all the experiments, with help from RO in strain construction and data analysis. NK hosted RKN his laboratory to perform the synthetic genetic array screens; NK also helped in proteomic analysis. RKN and JPC conceived the study, designed and interpreted the experiments, and wrote the manuscript.

## Supplementary Figure Legends

**Figure S1:**
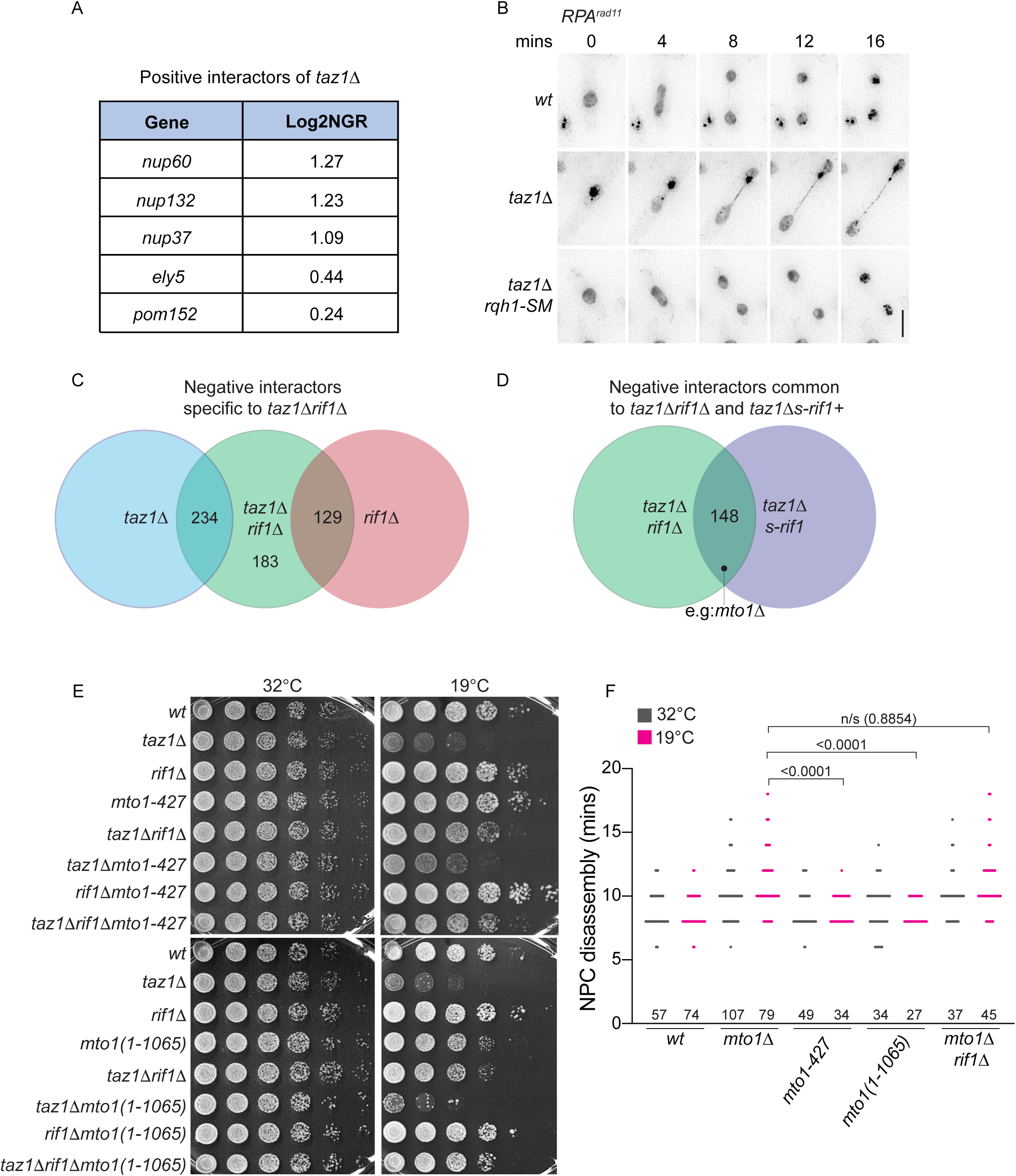
Characterization of cold-specific genetic interactors with *taz1Δ*. ***A***, List of NPC components that show cold-specific positive genetic interactions with *taz1+* deletion. ***B***, Representative images of mitotic cells that do or do not harbor aberrant RPA^Rad^^11^ bridges. The *rqh1-SM* allele suppresses the formation of aberrant RPA bridges in a *taz1Δ* setting. ***C***, Venn diagram of negative genetic interactions highlighting in the center the *taz1Δrif1Δ* double mutant-specific negative genetic interactors. ***D***, Venn diagram highlighting those negative genetic interactions that are common to *taz1Δrif1Δ* and *taz1ΔS-rif1+* strains; such interactions likely represent anaphase functions. ***E***, 5-fold serial dilutions were incubated at 32°C (2 days) or 19°C (7 days). ***F***, Quantitation of NPC disassembly time post anaphase onset of represented genotypes maintained in log phase at 32°C (1 day) or 19°C (3 days). For comparison, *wt* and *taz1Δ* data are replotted from Figure 3C. P values derived from Mann-Whitney test is represented above the brackets.

**Figure S2:**
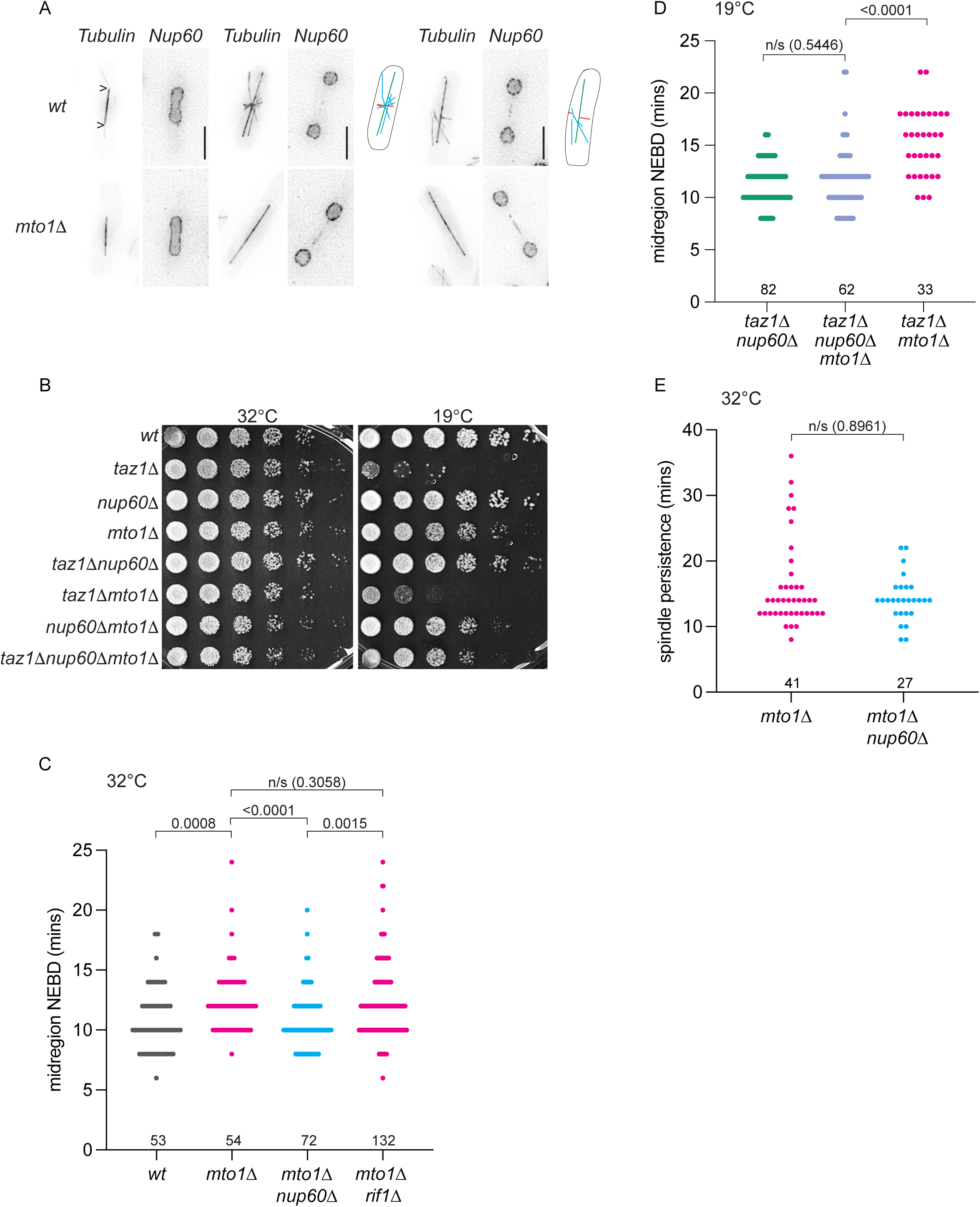
Delay in midregion NE breakdown, but not the spindle persistence, of *mto1Δ* cells inhibits telomere entanglement resolution. ***A***, Frames from films of mitotic cells with Nup60-mCherry and GFP-tubulin grown at 32°C. Arrowheads represent astral cytosolic microtubules nucleated from the SPB during mitosis. Cartoons representing the microtubules are shown to the right of each cell. Spindles are shown in green, equatorial microtubules in red and post anaphase arrays in blue. In *wt* cells equatorial and post anaphase arrays cross across the anaphase midregion. All the foregoing cytosolic microtubules types are absent in the absence of Mto1. ***B***, 5-fold serial dilutions were incubated at 32°C (2 days) or 19°C (7 days). ***C***, The timing of anaphase midregion NE breakdown is plotted as minutes post-anaphase onset as indicated by the loss of NLS-GFP-βGAL signal from the midregion in cells maintained in log-phase at 32°C (as in Figure 2F). ***D***, Timing of anaphase midregion NE breakdown of cells maintained in log phase for 3 days at 19°C (as in Figure 2G). For comparison, *taz1Δnup60Δ* data is re-plotted from Figure 2G. ***E***, Timing of spindle disassembly relative to anaphase onset in the represented genetic backgrounds. Cells were maintained in log-phase at 32°C. P value derived from Mann-Whitney test is represented above the brackets in C, D and E.

**Figure S3:**
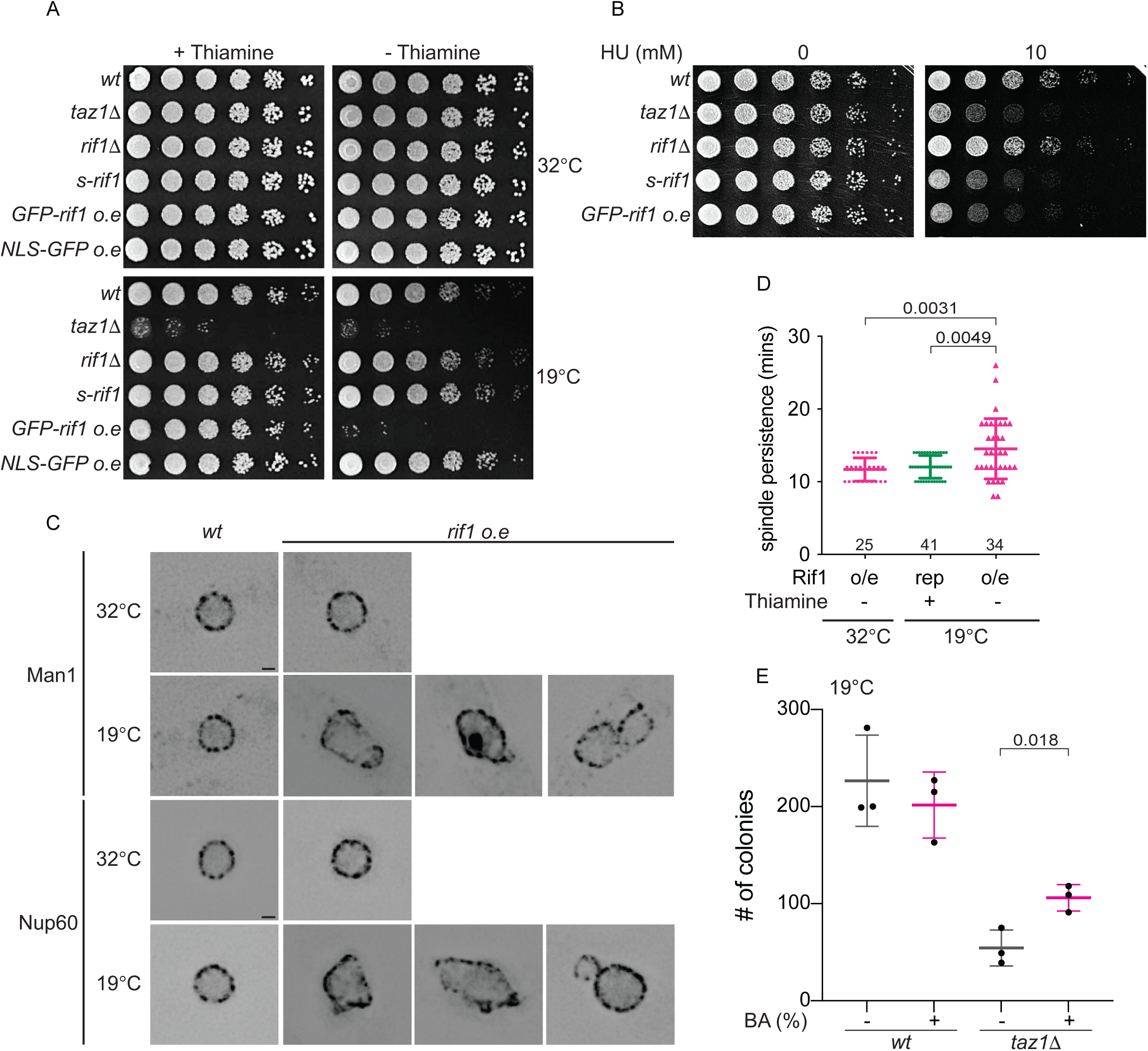
Rif1 overexpression causes cold-specific, lethal NE deformities. ***A***, 5-fold serial dilutions of cells grown to log phase in PMG minimal media with/without thiamine at 32°C were stamped on PMG-agar +/-thiamine and incubated at 32°C (2 days) or 19°C (10 days). ***B***, 5-fold serial dilutions of cells grown in PMG without thiamine to log phase at 32°C were stamped on plates lacking thiamine, with or without 10mM HU; the plates were incubated at 32°C for either 4 or 2 days, respectively. Cells overexpressing Rif1 experience replication stress. ***C***, Images of cells with or without overexpressed GFP-Rif1; the cells also expressed either Man1-tomato or Nup60-mCherry from their respective endogenous locus. Each image is a single Z slice with the maximum signal intensity. Scale bars represent 1μm. Rif1 overexpression causes cold-specific nuclear envelope irregularities. ***D***, Timing of spindle disassembly relative to anaphase onset. All the cells were maintained in log phase in PMG minimal media with (repressed) or without (de-repressed) thiamine. Delay in midregion NE breakdown causes spindle persistence in Rif1 overexpressing cells only when grown in the cold. P values from Mann-Whitney test is represented above the brackets. ***E***, Quantitation of number of colonies formed on plates at 32°C (without BA) from cultures grown with 0% or 0.075% BA at 19°C for 3 days. Each data point is an average of colonies formed from 3 technical replicates in which 300 cells were plated for each replicate. An average of three independent experiment is represented in the graph. P-value derived from two-tailed parametric t-test is represented over the brackets.

**Figure S4:**
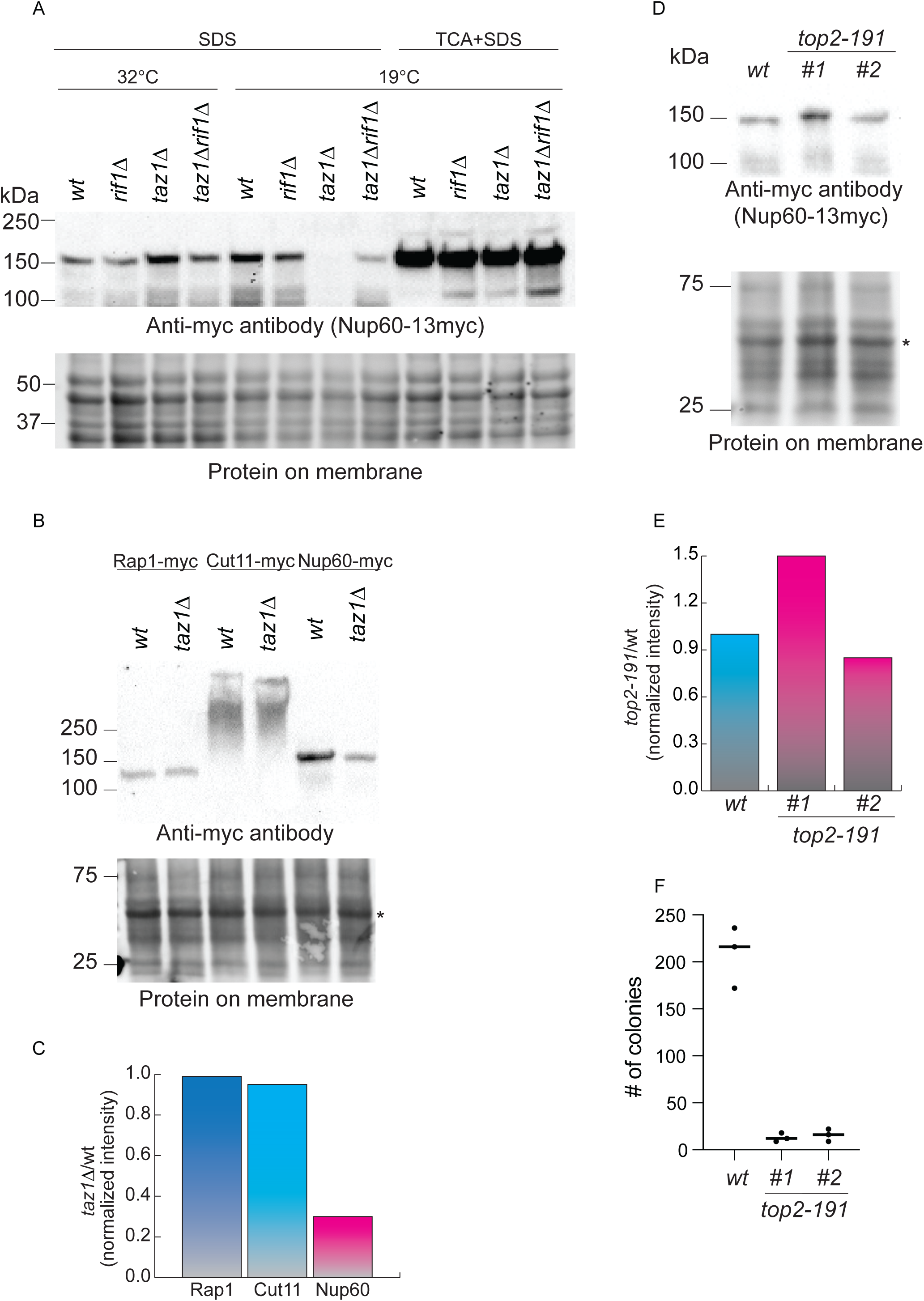
Rif1 alters the biochemistry of Nup60 extraction from *taz1Δ* cells in a cold-specific manner. ***A***, Western blot analysis of endogenously expressed Nup60-13myc from the cultures of the indicated genotypes. The total proteins were extracted in SDS buffer, from cells grown either at 32°C or 19°C, with or without TCA treatment. ***B***, Western blot analysis of represented proteins from cells in the presence or absence of Taz1 grown in cold for 3 days. ***C***, Quantitation of anti-myc signal from (B) normalized to the intensity of the band marked (*) on the lower panel of (B). ***D***, Western blot analysis of Nup60-13myc from cells with *wt* Top2 or temperature sensitive Top2 (Top2-191), grown at a non-permissive temperature (32°C) for 3 hours. Total protein was extracted in SDS buffer, without TCA treatment. #1 and #2 represent two different clones. ***E***, Quantitation of Nup60 intensity from the blot in (D), normalized to the intensity of the asterisked band. ***F***, Viability of colonies grown from the cultures used for western blot analysis in (D). 300 cells of each genotype were plated and incubated at permissive temperature (25°C) for 3 days. The low number of colonies formed in the *top2-191 nup60-13myc* cells represent loss in viability in culture grown at 32°C.

**Supplementary information for Figure S4: Rif1 alters the NE association of Nup60 in response to telomere entanglements** The epistatic interaction between *nup60*+ and *rif1*+ suggests that Nup60 and Rif1 act via the same pathway to control *taz1Δ* telomere detanglement. As we previously found that Rif1’s interaction with PP1 is required for inhibition of entanglement resolution^8^, our observation that Rif1 mediates cold specific NE changes hints that Rif1 may impact the phosphorylation status of Nup60, or of factors that affect Nup60 function, during telomere entanglement resolution.

To determine whether Nup60 is differentially modified in *taz1Δ* cells in a *rif1*+ dependent manner, we extracted endogenously tagged and functional Nup60-13Myc under standard denaturing conditions, 1% SDS (Figure S4A) or 8M Urea (data not shown). Surprisingly, while Nup60-13Myc is readily extractable from *wt* or *taz1Δ* cells grown at 32°C or *wt* cells grown at 19°C, extraction from *taz1Δ* cells grown at 19°C is profoundly inefficient (Figure S4A). This loss of recovery of Nup60 does not stem from lack of expression or mislocalization of Nup60 in *taz1Δ* cells in 19°C, as the Nup60-13Myc signal is readily seen by immunofluorescence (data not shown) and Nup60-mCherry shows peripheral nuclear localization similar to *wt* when viewed in live cells grown at 19°C (Figure 2B). Crucially, treating the cells with trichloroacetic acid (TCA) before extraction (Figure S4A), or direct extraction in guanidinium HCl buffer (data not shown), generally increases the extraction efficiency of Nup60-13Myc and allows full extraction of Nup60-13Myc from *taz1*Δ cells grown at 19°C. We considered that this might reflect an artifact due to upregulation in dying cells of detergent-resistant proteases that non-specifically degrade proteins upon cell rupture. However, the loss of Nup60 signal appears to be specific, as other subunits of the NPC (Cut11) or shelterin (Rap1) do not show similar behaviors (Figure S5C and D). Moreover, the loss of SDS-extractable Nup60 is not conserved in all dying cells, as the temperature-sensitive *top2-191* allele does not confer loss of Nup60 signal upon growth at nonpermissive temperature (Figure S4D-F).

The lack of Nup60 in SDS extracts of *taz1*Δ cells is Rif1-dependent, as removal of *rif1*+ restores SDS extraction from cells grown at 19°C (Figure S4A). Hence, Rif1-mediated modifications cause enhanced protease sensitivity and/or decreased extractability of Nup60 at 19°C. Whether Rif1 does this by modulating PP1 phosphatase activity on Nup60 or other proteins or phospholipids of the NE remains unknown. Indeed, we performed a phosphoproteomic comparison of *rif1+*, *rif1Δ* and *S-rif1+* cells grown at 32°C *versus* 19°C and found several Nup60 residues, most prominently S-157, that are hyperphosphorylated in a *rif1Δ/S-rif1* setting. However, mutation of S-157 has thus far yielded no consistent phenotype, which could mean that redundant phospho-sites on Nup60 compensate for loss of S-157, or that the relevant Rif1-modified protein is a Nup60-associated factor (or multiple factors) rather than solely Nup60. In either case, it is clear that Rif1 acts upstream of Nup60, altering its biochemistry, its extractability, and the propensity for NPC removal to allow anaphase midregion NE breakdown and entanglement resolution.

## Supplementary Tables

**Table S1:** Systematic IDs of genetic interactors from synthetic genetic array screens performed in the current study. Sheet1 and 2 provides the list of negative and positive interactors with *taz1Δ* at 19°C respectively. Sheet 3 provides a list of common negative interactors between *taz1*Δ*rif1*Δ and *taz1*Δ*S-rif1* at 19°C.

| *taz1Δ* negative interactors | *taz1Δ* positive interactors | Negative interactors common to *taz1Δrif1Δ* and *taz1ΔS-rif1+* (Gene name) |
| --- | --- | --- |
| SPAC2G11.10c | SPAC18B11.02c | SPAC1093.01 |
| SPAC1486.01 | SPAC1952.07 | SPAC110.02 |
| SPAC25A8.01c | SPBC1734.06 | SPAC1142.03c |
| SPAPB1A11.01 | SPCC70.03c | SPAC11E3.10 |
| SPBC16G5.13 | SPBC17G9.10 | SPAC13G6.10c |
| SPBC1604.20c | SPAC3A12.10 | SPAC1527.02 |
| SPBC30D10.13c | SPCC622.15c | SPAC1556.03 |
| SPBC16A3.01 | SPAC26H5.09c | SPAC1556.04c |
| SPAC1D4.01 | SPAC19A8.10 | SPAC17A2.09c |
| SPAC140.02 | SPAC17G6.17 | SPAC17A5.08 |
| SPAC637.09 | SPBC15D4.15 | SPAC17G8.13c |
| SPBC646.02 | SPAC16C9.05 | SPAC17H9.08 |
| SPAC17A5.02c | SPAC1B3.05 | SPAC1952.02 |
| SPBC3H7.12 | SPAC4G9.11c | SPAC19B12.06c |
| SPAC3F10.11c | SPAC694.06c | SPAC19G12.11 |
| SPCC757.09c | SPBC17D1.07c | SPAC1B3.16c |
| SPBC577.12 | SPAC869.06c | SPAC1D4.03c |
| SPCC74.09 | SPAC8C9.03 | SPAC20G8.10c |
| SPBC428.05c | SPBC609.05 | SPAC21E11.06 |
| SPCC1753.02c | SPAC20H4.10 | SPAC222.08c |
| SPAC21E11.03c | SPAC11H11.05c | SPAC22A12.02c |
| SPAC1F3.03 | SPBC1289.10c | SPAC22F8.09 |
| SPBC336.14c | SPCC23B6.03c | SPAC23G3.05c |
| SPAC20H4.09 | SPCC1020.11c | SPAC26A3.04 |
| SPAC5D6.04 | SPCP1E11.02 | SPAC27F1.08 |
| SPBC651.05c | SPBC29A3.10c | SPAC29B12.04 |
| SPAC23A1.09 | SPAC23H4.17c | SPAC29B12.14c |
| SPCC16C4.01 | SPAC23D3.09 | SPAC2F3.02 |
| SPAC8F11.10c | SPAC4H3.02c | SPAC2F3.07c |
| SPBC18H10.15 | SPAC1071.03c | SPAC2F7.09c |
| SPCC663.09c | SPBC23E6.02 | SPAC30D11.04c |
| SPAC8E11.06 | SPBC725.06c | SPAC30D11.06c |
| SPAC3C7.04 | SPBP8B7.13 | SPAC31G5.21 |
| SPBC1D7.04 | SPCC24B10.21 | SPAC3C7.13c |
| SPBC21D10.10 | SPBC839.03c | SPAC3G6.01 |
| SPAC8F11.03 | SPAC17A2.09c | SPAC3G6.05 |
| SPBC215.03c | SPBC56F2.08c | SPAC3G9.03 |
| SPBC1778.05c | SPBC29A10.05 | SPAC3G9.07c |
| SPAC1783.02c | SPBP4H10.18c | SPAC3H8.07c |
| SPBC1734.12c | SPCC306.04c | SPAC4D7.07c |
| SPCC5E4.10c | SPCC576.12c | SPAC4D7.11 |
| SPAC3G9.01 | SPAC343.11c | SPAC4G9.13c |
| SPCC162.12 | SPAC4G9.19 | SPAC4G9.16c |
| SPAC17G6.08 | SPAC13C5.04 | SPAC4G9.20c |
| SPAC12G12.01c | SPCC126.08c | SPAC513.06c |
| SPAC1F12.02c | SPBC16G5.02c | SPAC56F8.04c |
| SPBC31F10.13c | SPBC31F10.02 | SPAC56F8.09 |
| SPBC8E4.05c | SPCC1494.10 | SPAC56F8.15 |
| SPCC11E10.06c | SPBP4H10.17c | SPAC57A10.12c |
| SPAPYUG7.03c | SPCC1393.08 | SPAC644.11c |
| SPBC405.04c | SPBC1347.03 | SPAC644.13c |
| SPBC1778.02 (Rap1) | SPAC2C4.15c | SPAC688.10 |
| SPAC3A11.13 | SPBC32H8.03 | SPAC6B12.08 |
| SPCC23B6.05c | SPAC6B12.12 | SPAC6B12.16 |
| SPBC651.03c | SPAC13G7.06 | SPAC6F12.06 |
| SPAC6G9.16c | SPBC16E9.14c | SPAC6G9.01c |
| SPAC3H1.03 | SPAC17A5.07c (Upl2) | SPAC6G9.03c |
| SPCC736.07c | SPBC577.08c | SPAC6G9.09c |
| SPBC947.08c | SPBP8B7.22 | SPAC6G9.14 |
| SPCC417.02 | SPAC2F7.17 | SPAC7D4.02c |
| SPCC895.07 | SPAPB24D3.02c | SPAC7D4.04 |
| SPAC20H4.07 | SPBC1105.04c | SPAC821.05 |
| SPBC27.02c | SPAC9E9.13 | SPAC823.09c |
| SPAC8C9.16c | SPAC23H3.05c | SPAC823.10c |
| SPAC27E2.07 | SPBC887.04c | SPAC8C9.04 |
| SPAC1687.15 | SPBC25H2.03 | SPAC8C9.12c |
| SPAC1F8.04c | SPBC16H5.06 | SPAC8E11.10 |
| SPBC2F12.11c | SPCC364.03 | SPAC977.16c |
| SPAC17G8.14c | SPAPB21F2.02 | SPAC9E9.04 |
| SPCC645.06c | SPBC2D10.12 | SPAC9G1.03c |
| SPAC664.03 | SPCC285.13c (Nup60) | SPAPB1E7.06c |
| SPBC713.08 | SPAC513.06c | SPBC106.01 |
| SPAC1B1.04c | SPBC27B12.11c | SPBC106.03 |
| SPCC622.14 | SPBC1778.04 | SPBC12D12.07c |
| SPAC25H1.07 | SPAC4D7.03 | SPBC13E7.11 |
| SPCC548.07c | SPBP4H10.16c | SPBC13G1.10c |
| SPCC794.09c | SPAC1556.01c | SPBC14F5.07 |
| SPAC9G1.08c | SPBC83.09c | SPBC15D4.10c |
| SPCC550.14 | SPCC4B3.08 | SPBC1685.15c |
| SPAC4F10.19c | SPCC16A11.07 | SPBC16E9.11c |
| SPBC13E7.03c | SPBC902.02c | SPBC16G5.05c |
| SPCC1020.07 | SPBC1347.02 | SPBC1711.05 |
| SPAC1834.05 | SPAC2G11.13 | SPBC1773.08c |
| SPAC19G12.13c | SPAC926.07c | SPBC1861.07 |
| SPCC613.12c | SPAC29B12.05c | SPBC19C2.02 |
| SPAC16.01 | SPAC513.04 | SPBC1E8.03c |
| SPCC777.13 | SPBC2D10.11c | SPBC20F10.02c |
| SPBC18H10.07 | SPCC1235.03 | SPBC21B10.06c |
| SPCC1020.12c | SPBC2G5.01 | SPBC21C3.02c |
| SPBC16G5.14c | SPBC16A3.13 | SPBC21C3.17c |
| SPBC1604.03c | SPBC1734.15 | SPBC21D10.12 |
| SPAC328.06 | SPAC2F3.11 | SPBC25B2.07c |
| SPBC3H7.10 | SPAC1687.05 | SPBC27.06c |
| SPAC23H3.11c | SPAC20G4.04c | SPBC2G2.03c |
| SPBC337.03 | SPBC19F8.02 | SPBC32H8.09 |
| SPCC188.02 | SPBC16A3.10 | SPBC337.02c |
| SPAC9.02c | SPBC18H10.04c | SPBC337.07c |
| SPAC2F3.15 | SPBC365.06 | SPBC3H7.11 |
| SPAPB2B4.02 | SPAC3A11.02 | SPBC409.11 |
| SPBC24C6.10c | SPAC17C9.13c | SPBC428.04 |
| SPAPB1E7.11c | SPBC365.07c | SPBC428.14 |
| SPCC364.05 | SPAC6F6.13c | SPBC4B4.12c |
| SPCC663.12 | SPCC1840.12 | SPBC4F6.11c |
| SPCC1739.14 | SPAC1805.03c | SPBC530.01 |
| SPAC30D11.12 | SPAPB1A10.08 | SPBC577.11 |
| SPBC16C6.03c | SPAC57A10.09c | SPBC685.06 |
| SPAC17C9.08 | SPAC9E9.12c | SPBC776.14 |
| SPBC1348.13 | SPAC25B8.05 | SPBC83.02c |
| SPAC13G6.09 | SPBC3E7.11c | SPBC83.05 |
| SPAC1D4.09c | SPAC3C7.07c | SPBC83.17 |
| SPAC12G12.15 | SPBC8D2.02c | SPBC839.13c |
| SPCC4B3.15 | SPAC4G8.03c | SPBP22H7.04 |
| SPAC1805.07c | SPAC343.06c | SPBP35G2.02 |
| SPBC1921.03c | SPCC126.12 | SPBP8B7.10c |
| SPAC30.02c | SPAC1805.04 (Nup132) | SPBP8B7.22 |
| SPBC25B2.10 | SPBC31F10.15c | SPBPJ4664.01 |
| SPAC1B3.08 | SPAC664.02c | SPCC1183.07 |
| SPBC713.05 | SPBC2D10.06 | SPCC11E10.08 |
| SPAC1B3.03c | SPBC31E1.01c | SPCC1223.01 |
| SPBC17D11.04c | SPCC31H12.04c | SPCC1235.11 |
| SPAC31A2.02 | SPBC30D10.14 | SPCC126.02c |
| SPBC337.09 | SPAC12G12.07c | SPCC126.10 |
| SPCPJ732.02c | SPAC6B12.06c | SPCC132.03 |
| SPAC323.05c | SPBC2A9.04c | SPCC1393.03 |
| SPAC3H5.04 | SPBC2D10.17 | SPCC1450.07c |
| SPAC4F10.02 | SPBC660.12c | SPCC1672.03c |
| SPAC823.03 | SPBC106.20 | SPCC1739.10 |
| SPAC6F6.03c | SPBC1861.07 | SPCC1840.04 |
| SPCC622.16c | SPCC338.08 | SPCC1919.05 |
| SPAC31A2.12 | SPCC1223.04c | SPCC24B10.19c |
| SPAC57A7.08 | SPAC4F10.18 | SPCC285.16c |
| SPAC19D5.03 | SPCC594.04c | SPCC306.04c |
| SPBC1685.01 | SPAC13A11.06 | SPCC338.02 |
| SPAC1486.04c | SPBPB2B2.09c | SPCC417.07c (mto1) |
| SPBP35G2.09 | SPBC2D10.13 | SPCC576.01c |
| SPAC630.05 | SPBC25H2.15 | SPCC576.17c |
| SPBC14F5.10c | SPBPB10D8.06c | SPCC584.11c |
| SPCC1682.12c | SPAC23H4.08 | SPCC613.11c |
| SPBC428.08c | SPCC736.06 | SPCC63.06 |
| SPBC14F5.13c | SPBC1703.14c | SPCC663.13c |
| SPBC800.09 | SPACUNK4.15 | SPCC777.06c |
| SPBC21B10.13c | SPBP8B7.08c | SPCC962.04 |
| SPAC1071.02 | SPCC1442.05c | SPCC965.10 |
| SPAC2E1P5.03 | SPBC25H2.09 | SPCC965.11c |
| SPAC1F3.06c | SPCC4G3.05c | SPCC965.13 |
| SPCC584.02 | SPCC4B3.05c | SPCC965.14c |
| SPAC1039.04 | SPBC8D2.04 | SPCP1E11.05c |
| SPBC3B9.11c | SPBC29A3.14c | SPCP31B10.06 |
| SPBC215.14c | SPCPB16A4.06c |  |
| SPAC1851.02 | SPCC794.02 |  |
| SPAC10F6.11c | SPCC777.17c |  |
| SPAC3C7.03c | SPCC777.04 |  |
| SPBC13G1.08c | SPCC736.02 |  |
| SPCC63.08c | SPCC645.02 |  |
| SPAC637.10c | SPCC61.03 |  |
| SPBC359.03c | SPCC594.01 |  |
| SPAC29B12.11c | SPCC576.02 |  |
| SPAC29A4.09 | SPCC576.01c |  |
| SPBC947.15c | SPCC550.09 |  |
| SPBC3D6.13c | SPCC550.08 |  |
| SPBC354.09c | SPCC550.07 |  |
| SPAPB8E5.06c | SPCC4G3.11 |  |
| SPAC6C3.02c | SPCC4F11.03c |  |
| SPAC31G5.15 | SPCC4E9.01c |  |
| SPBC354.15 | SPCC417.06c |  |
| SPBC3B9.11c | SPCC364.06 |  |
| SPAC17C9.10 | SPCC338.18 |  |
| SPAC24H6.13 | SPCC320.03 |  |
| SPCP31B10.02 | SPCC306.10 |  |
| SPBC365.14c | SPCC2H8.05c |  |
| SPBC17G9.09 | SPCC24B10.19c |  |
| SPCC1223.05c | SPCC24B10.12 |  |
| SPAC1782.11 | SPCC1919.11 |  |
| SPBC8D2.17 | SPCC1919.09 |  |
| SPAC4D7.06c | SPCC191.10 |  |
| SPBC11C11.01 | SPCC18B5.10c |  |
| SPAPJ760.02c | SPCC18B5.09c |  |
| SPCC285.14 | SPCC18B5.05c |  |
| SPCC1919.13c | SPCC1840.07c |  |
| SPBC18E5.06 | SPCC1840.06 |  |
| SPAC1D4.11c | SPCC1795.03 |  |
| SPAC959.08 | SPCC1795.01c |  |
| SPCC162.08c | SPCC16C4.12 |  |
| SPBP35G2.07 | SPCC16A11.16c |  |
| SPBPB2B2.14c | SPCC1682.14 |  |
| SPBC36.04 | SPCC1672.12c |  |
| SPAC23H3.08c | SPCC1672.09 |  |
| SPCC4G3.17 | SPCC1620.08 |  |
| SPBC354.13 | SPCC1620.07c |  |
| SPACUNK4.13c | SPCC1620.04c |  |
| SPCP31B10.07 | SPCC1620.03 |  |
| SPCC188.10c | SPCC162.02c |  |
| SPBC557.04 | SPCC162.01c |  |
| SPAC22A12.14c | SPCC1450.16c |  |
| SPAP7G5.04c | SPCC1450.08c |  |
| SPAC17A2.11 | SPCC1450.06c |  |
| SPCC126.03 | SPCC1450.05c |  |
| SPAPB1A10.09 | SPCC1442.16c |  |
| SPAC6B12.02c | SPCC1442.14c |  |
| SPBP23A10.14c | SPCC1393.11 |  |
| SPAPB1E7.04c | SPCC1393.02c |  |
| SPAC630.09c | SPCC1322.03 |  |
| SPCC553.03 | SPCC1281.08 |  |
| SPBC29A10.14 | SPCC1235.05c |  |
| SPBC31F10.09c | SPCC1235.01 |  |
| SPAC19G12.06c | SPCC1223.13 |  |
| SPAC227.15 | SPCC1223.11 |  |
| SPBC3D6.05 | SPCC1223.03c |  |
| SPBC16G5.09 | SPCC11E10.07c |  |
| SPCPB16A4.03c | SPCC1020.06c |  |
| SPCC126.06 | SPBPB2B2.19c |  |
| SPCC663.06c | SPBPB2B2.18 |  |
| SPAC7D4.12c | SPBPB10D8.05c |  |
| SPAC15E1.06 | SPBPB10D8.02c |  |
| SPACUNK4.11c | SPBP8B7.26 |  |
| SPBC25B2.01 | SPBP8B7.25 |  |
| SPAPB24D3.06c | SPBP8B7.21 |  |
| SPAC1A6.09c | SPBP8B7.11 |  |
| SPCC1682.13 | SPBP8B7.06 |  |
| SPAC30D11.10 | SPBP8B7.02 |  |
| SPAC23H3.12c | SPBP4G3.03 |  |
| SPAC31G5.17c | SPBP35G2.12 |  |
| SPCC24B10.18 | SPBP35G2.03c |  |
| SPAC3H1.12c | SPBP23A10.05 |  |
| SPBC428.02c | SPBP22H7.04 |  |
| SPBC19C2.06c | SPBCPT2R1.01c |  |
| SPCC1235.15 | SPBC947.09 |  |
| SPBC119.06 | SPBC8E4.03 |  |
| SPBC2A9.13 | SPBC8D2.19 |  |
| SPBC30D10.16 | SPBC839.06 |  |
| SPCC1884.01 | SPBC800.07c |  |
| SPBC18H10.08c | SPBC800.02 |  |
| SPAC1399.01c | SPBC725.02 |  |
| SPCC306.07c | SPBC713.09 |  |
| SPAC750.01 | SPBC6B1.08c |  |
| SPAC17A5.01 | SPBC6B1.02 |  |
| SPCC970.07c | SPBC691.03c |  |
| SPAC3A11.11c | SPBC651.11c |  |
| SPBC660.10 | SPBC651.10 |  |
| SPBC26H8.05c | SPBC651.07 |  |
| SPAC3H8.05c | SPBC651.06 |  |
| SPBC28E12.02 | SPBC651.04 |  |
| SPAC3A12.06c | SPBC649.02 |  |
| SPAC23G3.12c | SPBC609.03 |  |
| SPAC17H9.06c | SPBC609.02 |  |
| SPCC24B10.06 | SPBC577.02 |  |
| SPBC8D2.10c | SPBC543.10 |  |
| SPAC5H10.10 | SPBC543.02c |  |
| SPAPB1A10.03 | SPBC530.05 |  |
| SPAC521.05 | SPBC530.04 |  |
| SPAC19A8.03 | SPBC4F6.16c |  |
| SPBC2D10.18 | SPBC4F6.09 |  |
| SPAC869.03c | SPBC4F6.08c |  |
| SPCC1020.09 | SPBC4F6.04 |  |
| SPCC61.05 | SPBC4C3.09 |  |
| SPBC25H2.16c | SPBC4C3.06 |  |
| SPAC25B8.11 | SPBC428.17c |  |
| SPBC1861.05 | SPBC428.10 |  |
| SPCC16A11.08 | SPBC428.03c |  |
| SPAC10F6.13c | SPBC409.10 |  |
| SPBC31A8.01c | SPBC405.06 |  |
| SPAC3F10.05c | SPBC405.01 |  |
| SPBC543.03c | SPBC4.06 |  |
| SPCC24B10.13 | SPBC3H7.05c |  |
| SPBC119.03 | SPBC3E7.16c |  |
| SPAPB24D3.07c | SPBC3E7.08c |  |
| SPBC713.11c | SPBC3E7.05c |  |
| SPBC119.14 | SPBC3B8.08 |  |
| SPAC17G6.03 | SPBC359.04c |  |
| SPAC56F8.06c | SPBC354.07c |  |
| SPAC977.05c | SPBC342.04 |  |
| SPBC685.04c | SPBC342.01c |  |
| SPBC16G5.06 | SPBC336.05c |  |
| SPAC1D4.06c | SPBC32H8.09 |  |
| SPBC1709.11c | SPBC32H8.02c |  |
| SPAC2E1P5.02c | SPBC32F12.12c |  |
| SPAC17C9.12 | SPBC31F10.17c |  |
| SPBC215.02 | SPBC31F10.08 |  |
| SPAC2F7.08c | SPBC31F10.03 |  |
| SPCC1795.10c | SPBC31E1.02c |  |
| SPAC1610.02c | SPBC317.01 |  |
| SPCC1322.15 | SPBC30D10.18c |  |
| SPBC16D10.11c | SPBC30D10.09c |  |
| SPBC1706.01 | SPBC30D10.05c |  |
| SPAC29B12.10c | SPBC2G5.03 |  |
| SPBC4F6.12 | SPBC2G2.17c |  |
| SPBC20F10.05 | SPBC2G2.06c |  |
| SPCC1682.16 | SPBC2G2.02 |  |
| SPAC15E1.09 | SPBC2F12.04 |  |
| SPAC17H9.10c | SPBC2F12.03c |  |
| SPAC16E8.01 | SPBC2D10.14c |  |
| SPCC364.02c | SPBC2D10.10c |  |
| SPAC30C2.04 | SPBC2D10.07c |  |
| SPBC19G7.09 | SPBC2D10.04 |  |
| SPAC20H4.05c | SPBC2D10.03c |  |
| SPAC823.14 | SPBC2A9.08c |  |
| SPBC1271.05c | SPBC2A9.07c |  |
| SPBC646.13 | SPBC2A9.03 |  |
| SPBC16E9.12c | SPBC29A3.09c |  |
| SPCC285.09c | SPBC29A10.11c |  |
| SPAC24B11.12c | SPBC29A10.07 |  |
| SPBP8B7.09c | SPBC29A10.06c |  |
| SPCC18.10 | SPBC28F2.11 |  |
| SPAC4A8.14 | SPBC28E12.04 |  |
| SPBC2G2.14 | SPBC26H8.03 |  |
| SPAC14C4.07 | SPBC25H2.14 |  |
| SPAC26A3.09c | SPBC25D12.05 |  |
| SPAC4H3.06 | SPBC25B2.11 |  |
| SPAC3G6.04 | SPBC25B2.04c |  |
| SPAC26F1.02 | SPBC25B2.03 |  |
| SPCC1840.09 | SPBC24C6.04 |  |
| SPBC29B5.04c | SPBC23G7.14 |  |
| SPCC1393.07c | SPBC23E6.03c |  |
| SPBPB7E8.01 | SPBC23E6.01c |  |
| SPCC622.01c | SPBC21D10.12 |  |
| SPAC1071.04c | SPBC21D10.11c |  |
| SPAC3C7.06c | SPBC21C3.09c |  |
| SPAC3F10.15c | SPBC21C3.08c |  |
| SPCC1259.11c | SPBC21C3.06 |  |
| SPAC25B8.03 | SPBC21B10.08c |  |
| SPCC63.13 | SPBC21B10.04c |  |
| SPAC227.11c | SPBC21B10.03c |  |
| SPAC9.13c | SPBC216.04c |  |
| SPBC16G5.07c | SPBC215.05 |  |
| SPBC2F12.05c | SPBC20F10.10 |  |
| SPBC21D10.09c | SPBC19G7.04 |  |
| SPAC26A3.16 | SPBC19F8.08 |  |
| SPAC1610.01 | SPBC19F8.06c |  |
| SPAC4H3.12c | SPBC19F8.03c |  |
| SPBC409.20c | SPBC19C7.12c |  |
| SPBC32H8.06 | SPBC19C7.10 |  |
| SPCC1322.11 | SPBC19C7.01 |  |
| SPAPB8E5.05 | SPBC1921.01c |  |
| SPACUNK4.17 | SPBC18H10.13 |  |
| SPAC57A10.10c | SPBC1861.06c |  |
| SPCC1672.06c | SPBC17G9.12c |  |
| SPAC24C9.02c | SPBC17G9.05 |  |
| SPBPJ4664.03 | SPBC17D11.08 |  |
| SPBC530.14c | SPBC17D11.03c |  |
| SPCC188.09c | SPBC17D1.05 |  |
| SPBC56F2.11 | SPBC17A3.06 |  |
| SPAC14C4.15c | SPBC1778.10c |  |
| SPBC29A3.05 | SPBC1773.06c |  |
| SPAC1805.05 | SPBC1773.02c |  |
| SPAC17G6.05c | SPBC1718.02 |  |
| SPBC365.20c | SPBC1711.08 |  |
| SPCC1322.14c | SPBC1703.03c |  |
| SPBC29A10.10c | SPBC16D10.03 |  |
| SPBC1709.04c | SPBC16C6.05 |  |
| SPAC105.02c | SPBC1685.02c |  |
| SPBC9B6.11c | SPBC1683.13c |  |
| SPAC1B1.03c | SPBC1683.08 |  |
| SPAC4C5.03 | SPBC1683.06c |  |
| SPCC594.05c | SPBC1604.19c |  |
| SPAC227.18 | SPBC1604.08c |  |
| SPAC17A2.13c | SPBC15C4.06c |  |
| SPAC57A10.03 | SPBC13E7.11 |  |
| SPAC29A4.06c | SPBC13E7.08c |  |
| SPCC1739.06c | SPBC13E7.07 |  |
| SPCC569.04 | SPBC13E7.06 |  |
| SPBC2F12.15c | SPBC1348.07 |  |
| SPAC1834.04 | SPBC1347.13c |  |
| SPBC19C2.09 | SPBC12C2.08 |  |
| SPAC23A1.11 | SPBC12C2.04 |  |
| SPAC3C7.12 | SPBC1289.14 |  |
| SPBC887.17 | SPBC1289.09 |  |
| SPBC146.04 | SPBC1289.06c |  |
| SPAC1565.06c | SPBC1271.06c |  |
| SPBC1348.03 | SPBC1271.01c |  |
| SPAC1556.06 | SPBC11G11.03 |  |
| SPAC27E2.02 | SPBC11C11.10 |  |
| SPBC146.10 | SPBC1198.07c |  |
| SPBC543.08 | SPBC1198.03c |  |
| SPBC106.12c | SPBC1198.01 |  |
| SPAC17H9.13c | SPBC119.08 |  |
| SPCC285.17 | SPBC1105.11c |  |
| SPAC1952.06c | SPBC1105.09 |  |
| SPCC1919.01 | SPAPYUK71.03c |  |
| SPAC17A2.01 | SPAPJ691.03 |  |
| SPAC17A2.10c | SPAPB8E5.04c |  |
| SPBC1921.07c | SPAPB2C8.01 |  |
| SPBC1A4.05 | SPAPB24D3.05c |  |
| SPBC543.05c | SPAPB24D3.04c |  |
| SPBC18H10.21c | SPAPB1A10.15 |  |
| SPCC16C4.09 | SPAP14E8.04 |  |
| SPBC4F6.10 | SPAP14E8.02 |  |
| SPAC22E12.03c | SPAC9G1.10c |  |
| SPAC17A5.16 | SPAC9G1.07 |  |
| SPAC1952.11c | SPAC9G1.06c |  |
| SPBC17G9.08c | SPAC9G1.05 |  |
| SPCC550.03c | SPAC9G1.02 |  |
| SPAC20G4.02c | SPAC9E9.14 |  |
| SPCC736.05 | SPAC977.14c |  |
| SPAC16A10.01 | SPAC926.06c |  |
| SPCC794.01c | SPAC926.02 |  |
| SPCC550.12 | SPAC9.12c |  |
| SPBC409.19c | SPAC9.10 |  |
| SPBC16G5.17 | SPAC8E11.10 |  |
| SPAC13G6.02c | SPAC869.11 |  |
| SPAC3G6.03c | SPAC824.09c |  |
| SPCC1919.12c | SPAC823.09c |  |
| SPBC1271.15c | SPAC821.07c |  |
| SPBC725.15 | SPAC7D4.02c |  |
| SPCC1393.13 | SPAC6F6.17 (rif1) |  |
| SPCC126.13c | SPAC6F12.15c |  |
| SPCC1739.01 | SPAC6C3.07 |  |
| SPBC11B10.02c | SPAC6B12.04c |  |
| SPBC1105.10 | SPAC694.04c |  |
| SPAC1805.12c | SPAC694.03 |  |
| SPBC16D10.07c | SPAC644.13c |  |
| SPAC3A11.09 | SPAC644.09 |  |
| SPBC12C2.03c | SPAC644.07 |  |
| SPAC644.08 | SPAC631.01c |  |
| SPBC18H10.11c | SPAC5H10.11 |  |
| SPBC1D7.03 | SPAC5H10.07 |  |
| SPBC3H7.06c | SPAC57A7.13 |  |
| SPBC18H10.18c | SPAC57A7.09 |  |
| SPAC2G11.04 | SPAC57A7.07c |  |
| SPAC15E1.10 | SPAC57A10.08c |  |
| SPAC222.12c | SPAC57A10.06 |  |
| SPBC21B10.12 | SPAC56F8.12 |  |
| SPAC750.06c | SPAC56F8.09 |  |
| SPAC25H1.02 | SPAC56F8.04c |  |
| SPBC29A3.12 | SPAC56E4.07 |  |
| SPAP8A3.07c | SPAC513.02 |  |
| SPAC6F6.01 | SPAC4H3.01 |  |
| SPBC1709.12 | SPAC4G9.16c |  |
| SPAC22E12.11c | SPAC4F8.10c |  |
| SPAC6G10.08 | SPAC4F10.16c |  |
| SPCC24B10.09 | SPAC4D7.11 |  |
| SPCC70.10 | SPAC4D7.01c |  |
| SPBC14F5.03c | SPAC4A8.10 |  |
| SPAC11D3.05 | SPAC4A8.02c |  |
| SPBC577.14c | SPAC3H5.09c |  |
| SPAC56F8.02 | SPAC3H5.07 |  |
| SPAC18G6.15 | SPAC3G9.03 |  |
| SPBC2G5.04c | SPAC3F10.09 |  |
| SPAC343.07 | SPAC3F10.02c |  |
| SPAC1783.08c | SPAC3C7.02c |  |
| SPCC1739.15 | SPAC3A12.17c |  |
| SPBC36.07 | SPAC32A11.02c |  |
| SPAC227.17c | SPAC323.04 |  |
| SPAC15A10.10 | SPAC31G5.11 |  |
| SPCC126.11c | SPAC30D11.11 |  |
| SPAC2C4.16c | SPAC30.01c |  |
| SPAC1B3.02c | SPAC2F7.10 |  |
| SPBC18E5.11c | SPAC2F7.09c |  |
| SPAC328.04 | SPAC2F7.04 |  |
| SPAC3A12.13c | SPAC2F7.02c |  |
| SPAC5D6.07c | SPAC2E12.03c |  |
| SPCPB16A4.05c | SPAC2C4.10c |  |
| SPAC8C9.11 | SPAC2C4.09 |  |
| SPBC365.04c | SPAC29B12.13 |  |
| SPAC26H5.05 | SPAC29B12.08 |  |
| SPCC663.17 | SPAC29B12.06c |  |
| SPAC25G10.04c | SPAC27F1.05c |  |
| SPCC790.02 | SPAC27D7.14c |  |
| SPCC622.19 | SPAC27D7.13c |  |
| SPBC660.17c | SPAC27D7.08c |  |
| SPCC1442.17c | SPAC27D7.04 |  |
| SPBC18H10.10c | SPAC25H1.09 |  |
| SPAC4G8.07c | SPAC25H1.06 |  |
| SPBC21B10.10 | SPAC25G10.05c |  |
| SPAPB2B4.06 | SPAC25B8.04c |  |
| SPBC21D10.07 | SPAC24H6.03 |  |
| SPAC1F5.08c | SPAC24B11.14 |  |
| SPBC2A9.11c | SPAC23H4.02 |  |
| SPBC16C6.08c | SPAC23H3.09c |  |
| SPBC19G7.16 | SPAC23G3.07c |  |
| SPCP20C8.03 | SPAC23G3.04 |  |
| SPAC4A8.03c | SPAC23D3.13c |  |
| SPBC36.06c | SPAC23C4.08 |  |
| SPBC800.08 | SPAC23C4.05c |  |
| SPAC4H3.07c | SPAC23C4.02 |  |
| SPAC977.02 | SPAC22H10.03c |  |
| SPBP23A10.10 | SPAC22H10.02 |  |
| SPCC1322.02 | SPAC22G7.08 |  |
| SPBC16H5.05c | SPAC22G7.03 |  |
| SPAC3H8.02 | SPAC22F8.03c |  |
| SPAC21E11.05c | SPAC22F3.13 |  |
| SPCC663.14c | SPAC22F3.06c |  |
| SPAC6G10.02c | SPAC22F3.03c |  |
| SPCC5E4.05c | SPAC22E12.19 |  |
| SPAC23A1.07 | SPAC22E12.14c |  |
| SPAC4G8.11c | SPAC22E12.04 |  |
| SPAC186.05c | SPAC227.14 |  |
| SPAC23C4.12 | SPAC227.03c |  |
| SPBC2D10.09 | SPAC222.05c |  |
| SPCC1906.02c | SPAC20H4.11c |  |
| SPBC337.16 | SPAC20G4.03c |  |
| SPAC22F3.12c | SPAC1F8.03c |  |
| SPAC31A2.16 | SPAC1F7.09c |  |
| SPBC609.04 | SPAC1F7.06 |  |
| SPBC4.01 | SPAC1B3.17 |  |
| SPBC4B4.04 | SPAC1B3.15c |  |
| SPAC13G7.13c | SPAC1B3.11c |  |
| SPAC3H8.04 | SPAC1B3.10c |  |
| SPBC16E9.02c | SPAC1A6.06c |  |
| SPAC3F10.07c | SPAC1A6.04c |  |
| SPAC6F12.09 | SPAC19G12.15c |  |
| SPBC12D12.06 | SPAC19D5.01 |  |
| SPBC3B9.05 | SPAC19B12.08 |  |
| SPBC115.03 | SPAC19B12.07c |  |
| SPBC660.14 | SPAC1952.09c |  |
| SPBC19C2.04c | SPAC1952.03 |  |
| SPBC216.01c | SPAC18G6.12c |  |
| SPAC227.10 | SPAC18B11.03c |  |
| SPCC18B5.06 | SPAC186.09 |  |
| SPAC922.04 | SPAC1834.09 |  |
| SPBC713.03 | SPAC17H9.19c |  |
| SPCP20C8.01c | SPAC17H9.14c |  |
| SPAC3H8.08c | SPAC17C9.11c |  |
| SPAC688.06c | SPAC17A5.04c |  |
| SPAC17C9.07 | SPAC1782.08c |  |
| SPBC83.18c | SPAC1782.05 |  |
| SPAC4F8.08 | SPAC16E8.09 |  |
| SPAPB1E7.08c | SPAC16C9.07 |  |
| SPBC1773.14 | SPAC16A10.07c |  |
| SPBC23G7.15c | SPAC1687.19c |  |
| SPBC30B4.02c | SPAC1687.10 |  |
| SPCC16C4.10 | SPAC167.07c |  |
| SPBC17D11.01 | SPAC1635.01 |  |
| SPAC24B11.10c | SPAC15E1.05c |  |
| SPAC513.07 | SPAC15A10.16 |  |
| SPCC16A11.15c | SPAC1556.04c |  |
| SPAC343.04c | SPAC1527.01 |  |
| SPAPB24D3.08c | SPAC144.11 |  |
| SPCC330.06c | SPAC13G7.03 |  |
| SPAC9E9.10c | SPAC1399.05c |  |
| SPCC4B3.04c | SPAC139.06 |  |
| SPAC821.09 | SPAC139.04c |  |
| SPAC19B12.11c | SPAC12G12.03 |  |
| SPAC12B10.03 | SPAC12B10.14c |  |
| SPBC16H5.04 | SPAC12B10.13 |  |
| SPBC1921.06c | SPAC12B10.09 |  |
| SPBC119.04 | SPAC12B10.01c |  |
| SPAPJ696.01c | SPAC1296.01c |  |
| SPBC3D6.06c | SPAC11H11.01 |  |
| SPBC23G7.13c | SPAC11E3.10 |  |
| SPAC631.02 | SPAC11E3.01c |  |
| SPCC1393.05 | SPAC11D3.08c |  |
| SPCC24B10.14c | SPAC110.02 |  |
| SPAC13G7.11 | SPAC10F6.16 |  |
| SPBC1198.14c | SPAC10F6.15 |  |
| SPBC215.07c | SPAC1071.06 |  |
| SPAC22F8.04 | SPAC1006.06 |  |
| SPAC664.04c | SPAC1002.17c |  |
| SPBC1773.01 |  |  |
| SPAC56E4.06c |  |  |
| SPBC24C6.08c |  |  |
| SPCC18.06c |  |  |
| SPAC3C7.10 |  |  |
| SPAC694.02 |  |  |
| SPBC106.13 |  |  |
| SPCC825.04c |  |  |
| SPBP23A10.16 |  |  |
| SPBC26H8.09c |  |  |
| SPBC725.04 |  |  |
| SPAC26A3.01 |  |  |
| SPBC119.16c |  |  |
| SPAC9G1.04 |  |  |
| SPAC13G7.02c |  |  |
| SPAC328.01c |  |  |
| SPBC3H7.13 |  |  |
| SPCC132.01c |  |  |
| SPAC22E12.01 |  |  |

**Table S2:**
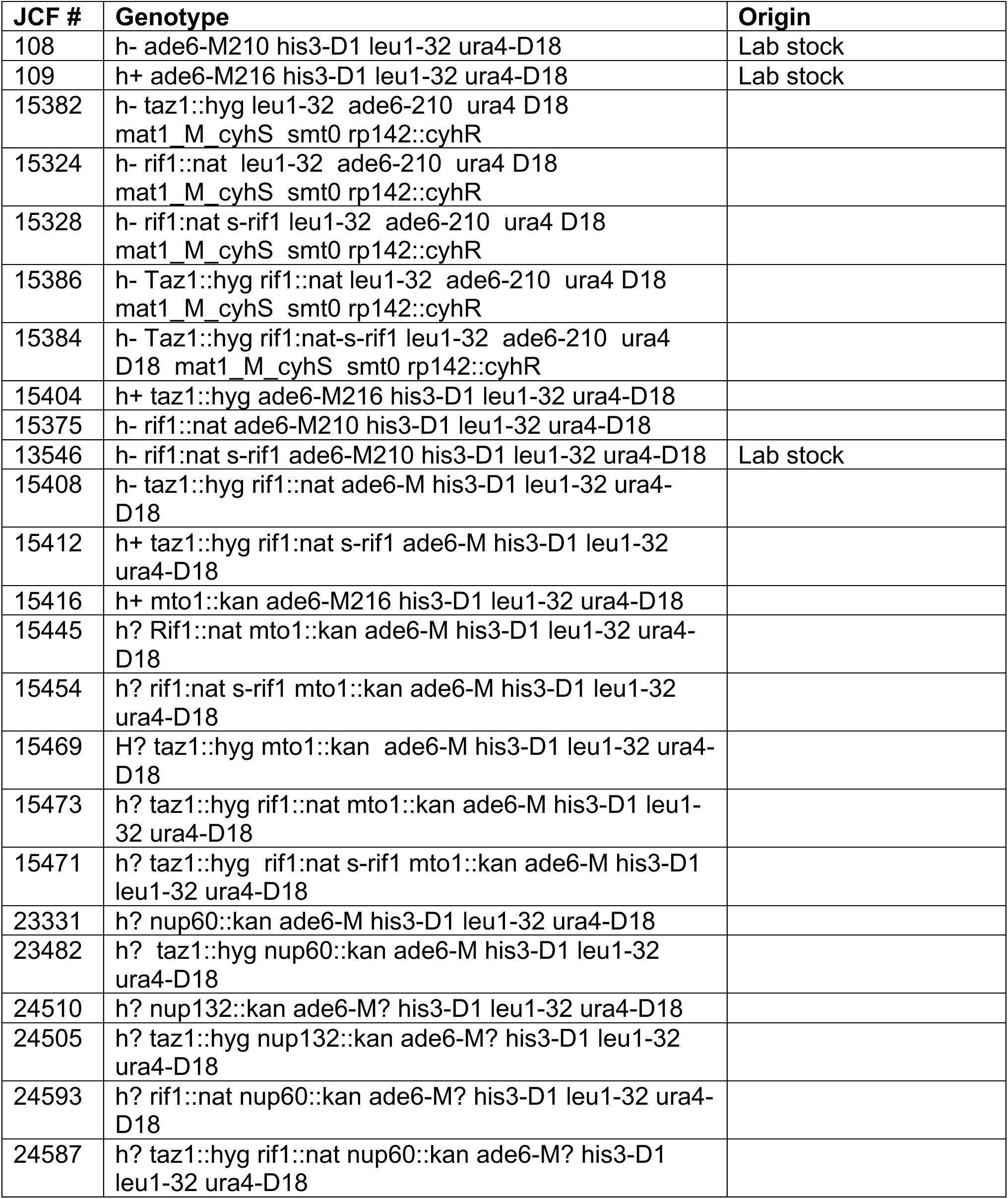
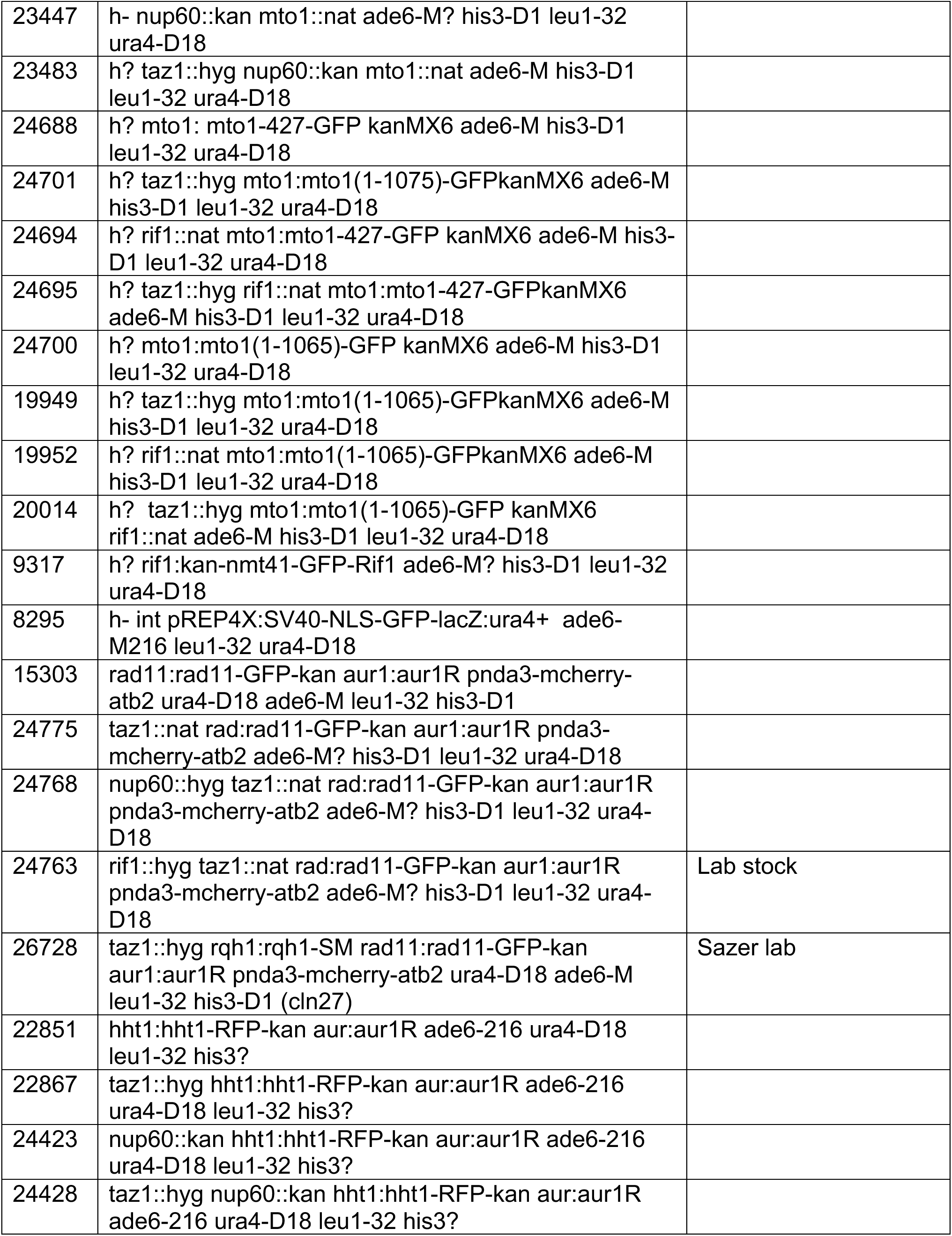
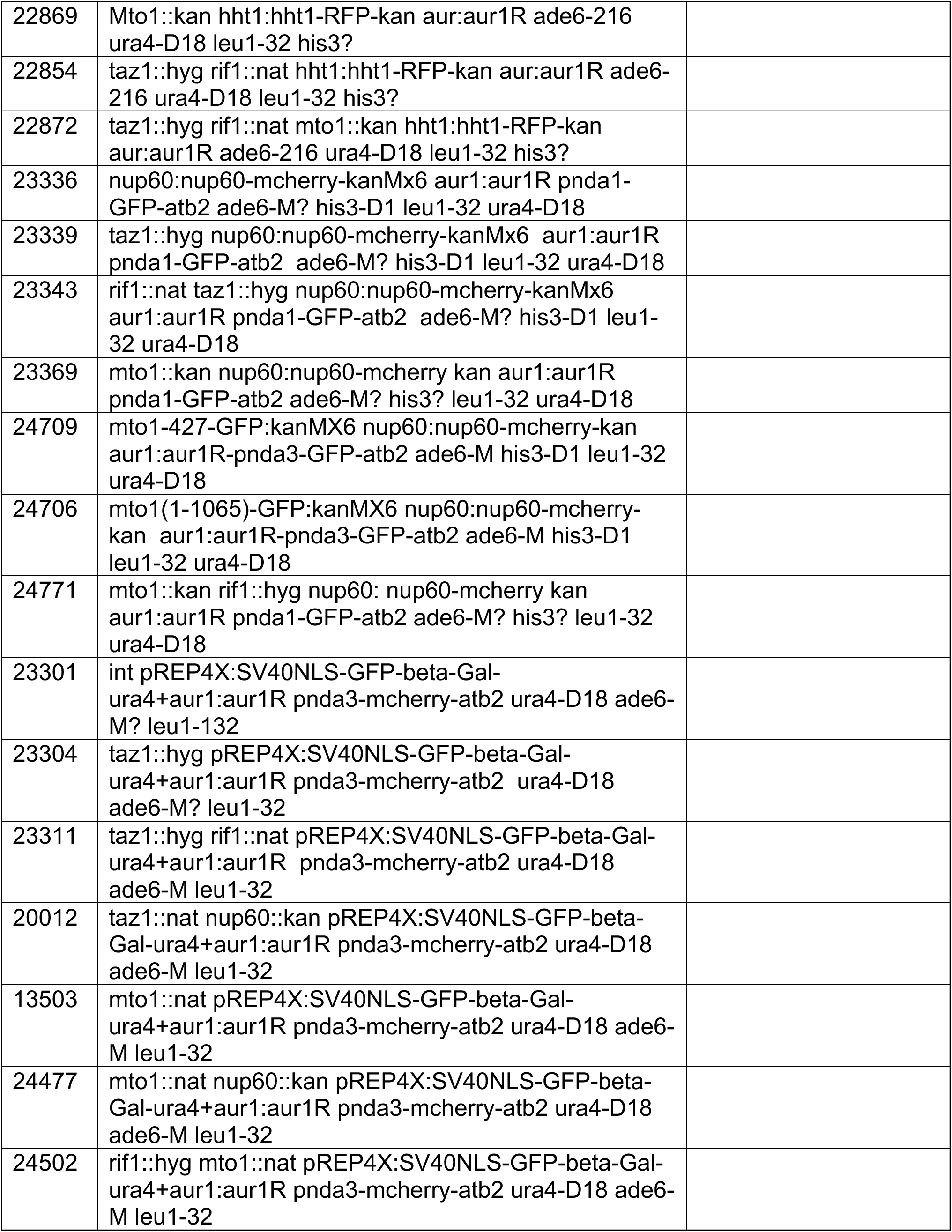
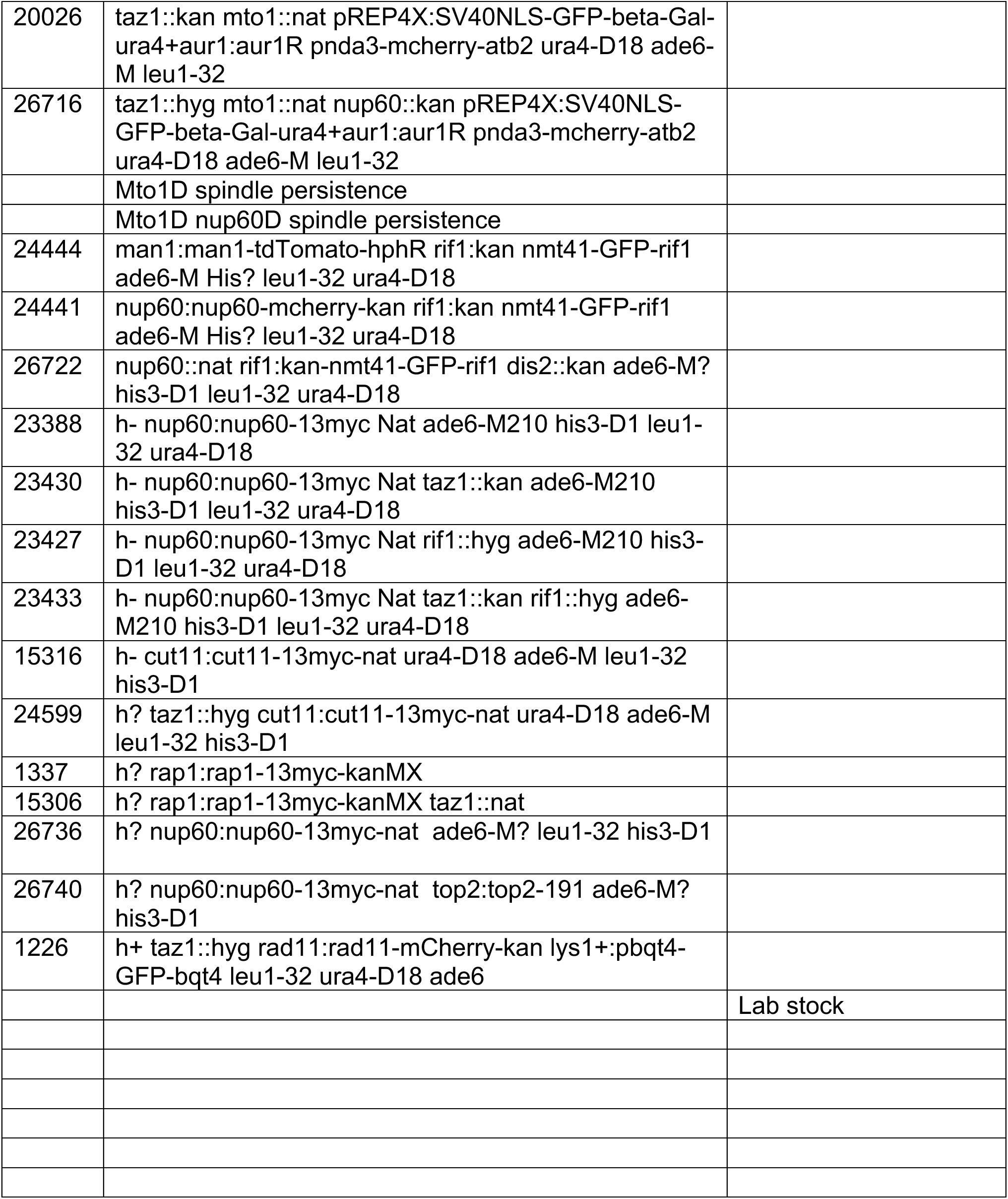
List of strains used in the current study.

**Supplementary Videos S1-4:** Time lapse videos of mitotically dividing *taz1Δ* cells maintianed at 32°C, expressing GFP-bqt4 (NE marker) and Rad11-mcherry (RPA). Also see Figure 3D.

## Methods

### Media

Media comprised YES broth (Sunrise #2011-500), YES agar (Sunrise #2012-500), PMG (Sunrise #2060-500). α-myc antibody was BioXCell – 9E10 used at 1:5000 dilution. Precast gels (Mini-PROTEAN TGX Stain-Free Gels #4568084) were used with Biorad buffer (#1610732 and #1610734).Media and growth conditions were as previously described^66^. Strains are listed in Table S1.

### Strain construction

Gene deletions and tag insertions were generated as described previously^67^ and used to create further strains through crossing, sporulation and selection. Tagging was further confirmed by sequencing the tag-gene junctions. Diploids with heterozygous gene deletions were created either by gene deletion in preselected diploids or crossing two haploids followed by diploid selection via *ade6-210/216* complementation.

### Synthetic genetic array screening

Single or double mutants of *taz1Δ/rif1Δ/S-rif1+* were generated in the PEM-2 strain background as described previously^68, 69^. The query strains were crossed with the Bioneer *S. pombe* gene deletion library. Double or triple mutant haploids were selected using the appropriate antibiotics. Colonies were plated robotically as described earlier^69^ with a few modifications, most prominently repetitive stamping (two additional rounds) of colonies still in log phase on YES plates after the final selection of progeny. This procedure was developed to ensure that we screen for c/s rather than for recovery from stationary phase at cold temperature. The final stamping was performed in duplicate, with one incubated at 30°C and the other at 19°C. After 3 days of growth the plates were imaged, and growth (colony size) was measured as described earlier^69^. Growth of each genotype was normalized to the median growth of a given plate. The normalized growth ratio (19°C/30°C) was calculated for each genotype as a measure of genetic interaction. The outliers were determined by the 2-standard deviation method.

### Cell growth

All experiments were performed on logarithmically growing cells, which were grown in either YES or PMG with supplements, from a single colony at the indicated temperature. The log phase cells (0.5 OD) were diluted to 0.08-0.1 OD, incubated at 19°C and maintained in log phase by dilution with fresh media every day. This approach was used for microscopy and western blot analysis.

### Benzyl alcohol (BA) treatment

Cultures were grown as described above to saturation, diluted to 0.1 OD in YES broth without BA and incubated at 32°C. Once the cultures reached OD 0.5, they were diluted to 0.1 OD in YES broth with varying concentrations of BA (0, 0.0025%, 0.05 or 0.075%) and transferred to 19°C. The media was replaced daily with fresh broth containing the respective concentrations of BA. After 3 days, the cells were stained with Erythrosine B, imaged and counted using a hemocytometer; 300 cells were plated on YES media without BA and incubated at 32°C for 2 days.

### Microscopy

Mid log-phase cells were adhered to 35-mm glass culture dishes (MatTek Corporation) using 0.2 mg/ml of soybean lectin (Sigma-Aldrich) and filled with PMG broth. Time lapse images were captured in an environmental chamber set at 32°C with a DeltaVision Ultra with a 60x/1.42 (lens ID 10612) objective with 1.585 oil immersion, and a charge coupled device EDGE sCMOS camera. Cells were imaged every 2 mins for 40 minutes over 16 focal planes of 0.4 μm. These acquired images were deconvolved (conserved ratio method) and quick-projected into a 2D image using the maximum intensity setting in SoftWorx (Applied precision). Images using the DeltaVision OMX Flex (at conventional mode) were acquired and processed similarly.

### Top2-191 experiment

Pre-cultures were inoculated from a single colony grown overnight at 25°C, a permissive temperature for *top2-191*. The cells were diluted to 0.1 OD, grown to 0.5 OD at 25°C, diluted to 0.2 OD and transferred to 32°C for 3 hours. The cells were monitored for degree of cell death every hour by light microscopy. After 3 hours the cells were harvested and protein was extracted as described below. The same cultures were used for plating assays as described for BA treatment except that the plates were incubated at 25°C for 3 days.

### Protein extraction and western blot

25 OD (50mL of 0.5 OD culture) of log phase cultures were pelleted and washed three times with autoclaved water before protein extraction. The cells were ruptured using a beadbeater with 1.9g of glass beads either in SDS buffer (1X phosphate buffered saline, 10% Glycerol, 2mM EDTA, 2mM DTT, Roche protease inhibitor cocktail, 2mM PMSF, 1%SDS), 8M urea buffer, or 6M guanidinium HCl^70^. As incidated, cells were treated with 20% trichloroacetic acid and neutralized with 1M tris base before harvesting in SDS buffer.

For western blot analysis, 100 μg total protein was resolved on a BioRad Stainfree precast gel and transferred onto a PVDF membrane using tris-Glycine buffer with 20% methanol. The membrane was blocked with 5% non-fat milk and washed with PBS containing 0.05% Tween 20. Membranes were incubated in primary antibody (in blocking buffer) followed by HRP conjugated secondary antibody (in PBST), and developed in a BioRad Imager using ECL reagent (Amersham; RPN2236).

### Spindle analysis

Mitotic cell progression was monitored by observing spindle dynamics. In an unperturbed cell the metaphase spindle is maintained at <3μm length and elongates exponentially during anaphase.

### Statistical analysis

Statistical tests were performed in GrapPad Prism. Non-parametric analysis using the Mann-Whitney test method was performed to analyze anaphase midregion NE breakdown, NPC disassembly, % dead cells, viability, and spindle persistence experiments. A two tailed Fisher’s exact T test was performed to determine significance when analyzing percent RPA bridges and histone segregation. Double blind experiments were performed for Figure S4C. Two independent biological clones were analyzed for most experiments. No specific statistical methods were employed to predetermine *n*-values.

## References

1. Guttinger, S., Laurell, E. & Kutay, U. Orchestrating nuclear envelope disassembly and reassembly during mitosis. Nat Rev Mol Cell Biol 10, 178–191 (2009).

2. Friederichs, J.M. et al. The SUN protein Mps3 is required for spindle pole body insertion into the nuclear membrane and nuclear envelope homeostasis. PLoS Genet 7, e1002365 (2011).

3. Jaspersen, S.L. & Ghosh, S. Nuclear envelope insertion of spindle pole bodies and nuclear pore complexes. Nucleus 3, 226–236 (2012).

4. Smoyer, C.J. & Jaspersen, S.L. Breaking down the wall: the nuclear envelope during mitosis. Curr Opin Cell Biol 26, 1–9 (2014).

5. Fernandez-Alvarez, A., Bez, C., O’Toole, E.T., Morphew, M. & Cooper, J.P. Mitotic Nuclear Envelope Breakdown and Spindle Nucleation Are Controlled by Interphase Contacts between Centromeres and the Nuclear Envelope. Dev Cell 39, 544–559 (2016).

6. Dey, G. et al. Closed mitosis requires local disassembly of the nuclear envelope. Nature 585, 119–123 (2020).

7. Exposito-Serrano, M., Sanchez-Molina, A., Gallardo, P., Salas-Pino, S. & Daga, R.R. Selective Nuclear Pore Complex Removal Drives Nuclear Envelope Division in Fission Yeast. Curr Biol 30, 3212–3222 e3212 (2020).

8. Zaaijer, S., Shaikh, N., Nageshan, R.K. & Cooper, J.P. Rif1 Regulates the Fate of DNA Entanglements during Mitosis. Cell Rep 16, 148–160 (2016).

9. Chan, K.L. & Hickson, I.D. New insights into the formation and resolution of ultra-fine anaphase bridges. Semin Cell Dev Biol 22, 906–912 (2011).

10. Chan, K.L., Palmai-Pallag, T., Ying, S. & Hickson, I.D. Replication stress induces sister-chromatid bridging at fragile site loci in mitosis. Nat Cell Biol 11, 753–760 (2009).

11. Ait Saada, A., et al. Unprotected Replication Forks Are Converted into Mitotic Sister Chromatid Bridges. Mol Cell 66, 398–410 e394 (2017).

12. Chan, K.L., North, P.S. & Hickson, I.D. BLM is required for faithful chromosome segregation and its localization defines a class of ultrafine anaphase bridges. EMBO J 26, 3397–3409 (2007).

13. Sofueva, S. et al. Ultrafine anaphase bridges, broken DNA and illegitimate recombination induced by a replication fork barrier. Nucleic Acids Res 39, 6568–6584 (2011).

14. Baumann, C., Korner, R., Hofmann, K. & Nigg, E.A. PICH, a centromere-associated SNF2 family ATPase, is regulated by Plk1 and required for the spindle checkpoint. Cell 128, 101–114 (2007).

15. Hengeveld, R.C. et al. Rif1 Is Required for Resolution of Ultrafine DNA Bridges in Anaphase to Ensure Genomic Stability. Dev Cell 34, 466–474 (2015).

16. de Lange, T. Shelterin-Mediated Telomere Protection. Annu Rev Genet 52, 223–247 (2018).

17. Hou, H. & Cooper, J.P. Stretching, scrambling, piercing and entangling: Challenges for telomeres in mitotic and meiotic chromosome segregation. Differentiation 100, 12–20 (2018).

18. Cooper, J.P., Nimmo, E.R., Allshire, R.C. & Cech, T.R. Regulation of telomere length and function by a Myb-domain protein in fission yeast. Nature 385, 744–747 (1997).

19. Nimmo, E.R., Pidoux, A.L., Perry, P.E. & Allshire, R.C. Defective meiosis in telomere-silencing mutants of Schizosaccharomyces pombe. Nature 392, 825–828 (1998).

20. Ferreira, M.G. & Cooper, J.P. Two modes of DNA double-strand break repair are reciprocally regulated through the fission yeast cell cycle. Genes Dev 18, 2249–2254 (2004).

21. Ferreira, M.G. & Cooper, J.P. The fission yeast Taz1 protein protects chromosomes from Ku-dependent end-to-end fusions. Mol Cell 7, 55–63 (2001).

22. Miller, K.M., Rog, O. & Cooper, J.P. Semi-conservative DNA replication through telomeres requires Taz1. Nature 440, 824–828 (2006).

23. Dehe, P.M., Rog, O., Ferreira, M.G., Greenwood, J. & Cooper, J.P. Taz1 enforces cell-cycle regulation of telomere synthesis. Mol Cell 46, 797–808 (2012).

24. Sfeir, A. et al. Mammalian telomeres resemble fragile sites and require TRF1 for efficient replication. Cell 138, 90–103 (2009).

25. Zimmermann, M., Kibe, T., Kabir, S. & de Lange, T. TRF1 negotiates TTAGGG repeat-associated replication problems by recruiting the BLM helicase and the TPP1/POT1 repressor of ATR signaling. Genes Dev 28, 2477–2491 (2014).

26. Yang, Z., Takai, K.K., Lovejoy, C.A. & de Lange, T. Break-induced replication promotes fragile telomere formation. Genes Dev 34, 1392–1405 (2020).

27. Yang, Z., Sharma, K. & de Lange, T. TRF1 uses a noncanonical function of TFIIH to promote telomere replication. Genes Dev 36, 956–969 (2022).

28. Rog, O., Miller, K.M., Ferreira, M.G. & Cooper, J.P. Sumoylation of RecQ helicase controls the fate of dysfunctional telomeres. Mol Cell 33, 559–569 (2009).

29. Miller, K.M. & Cooper, J.P. The telomere protein Taz1 is required to prevent and repair genomic DNA breaks. Mol Cell 11, 303–313 (2003).

30. Miller, K.M., Ferreira, M.G. & Cooper, J.P. Taz1, Rap1 and Rif1 act both interdependently and independently to maintain telomeres. EMBO J 24, 3128–3135 (2005).

31. Hardy, C.F., Sussel, L. & Shore, D. A RAP1-interacting protein involved in transcriptional silencing and telomere length regulation. Genes Dev 6, 801–814 (1992).

32. Kanoh, J. & Ishikawa, F. spRap1 and spRif1, recruited to telomeres by Taz1, are essential for telomere function in fission yeast. Curr Biol 11, 1624–1630 (2001).

33. Hayano, M. et al. Rif1 is a global regulator of timing of replication origin firing in fission yeast. Genes Dev 26, 137–150 (2012).

34. Yamazaki, S. et al. Rif1 regulates the replication timing domains on the human genome. EMBO J 31, 3667–3677 (2012).

35. Kanoh, Y. et al. Rif1 binds to G quadruplexes and suppresses replication over long distances. Nat Struct Mol Biol 22, 889–897 (2015).

36. Dave, A., Cooley, C., Garg, M. & Bianchi, A. Protein phosphatase 1 recruitment by Rif1 regulates DNA replication origin firing by counteracting DDK activity. Cell Rep 7, 53–61 (2014).

37. Mattarocci, S. et al. Rif1 controls DNA replication timing in yeast through the PP1 phosphatase Glc7. Cell Rep 7, 62–69 (2014).

38. Cornacchia, D. et al. Mouse Rif1 is a key regulator of the replication-timing programme in mammalian cells. EMBO J 31, 3678–3690 (2012).

39. Hiraga, S. et al. Rif1 controls DNA replication by directing Protein Phosphatase 1 to reverse Cdc7-mediated phosphorylation of the MCM complex. Genes Dev 28, 372–383 (2014).

40. Hiraga, S.I. et al. Human RIF1 and protein phosphatase 1 stimulate DNA replication origin licensing but suppress origin activation. EMBO Rep 18, 403–419 (2017).

41. Hiraga, S.I. et al. Budding yeast Rif1 binds to replication origins and protects DNA at blocked replication forks. EMBO Rep 19 (2018).

42. Sawin, K.E., Lourenco, P.C. & Snaith, H.A. Microtubule nucleation at non-spindle pole body microtubule-organizing centers requires fission yeast centrosomin-related protein mod20p. Curr Biol 14, 763–775 (2004).

43. Venkatram, S. et al. Identification and characterization of two novel proteins affecting fission yeast gamma-tubulin complex function. Mol Biol Cell 15, 2287–2301 (2004).

44. Samejima, I., Miller, V.J., Rincon, S.A. & Sawin, K.E. Fission yeast Mto1 regulates diversity of cytoplasmic microtubule organizing centers. Curr Biol 20, 1959–1965 (2010).

45. Kanoh, Y., Ueno, M., Hayano, M., Kudo, S. & Masai, H. Aberrant association of chromatin with nuclear periphery induced by Rif1 leads to mitotic defect. Life Sci Alliance 6 (2023).

46. Kramarz, K. et al. The nuclear pore primes recombination-dependent DNA synthesis at arrested forks by promoting SUMO removal. Nat Commun 11, 5643 (2020).

47. Su, X.A., Dion, V., Gasser, S.M. & Freudenreich, C.H. Regulation of recombination at yeast nuclear pores controls repair and triplet repeat stability. Genes Dev 29, 1006–1017 (2015).

48. Pinzaru, A.M. et al. Replication stress conferred by POT1 dysfunction promotes telomere relocalization to the nuclear pore. Genes Dev 34, 1619–1636 (2020).

49. Ebrahimi, H., Masuda, H., Jain, D. & Cooper, J.P. Distinct ‘safe zones’ at the nuclear envelope ensure robust replication of heterochromatic chromosome regions. Elife 7 (2018).

50. Mattarocci, S. et al. Rif1 maintains telomeres and mediates DNA repair by encasing DNA ends. Nat Struct Mol Biol 24, 588–595 (2017).

51. Masai, H. et al. Rif1 promotes association of G-quadruplex (G4) by its specific G4 binding and oligomerization activities. Sci Rep 9, 8618 (2019).

52. Fox, C.A. & Gartenberg, M.R. Palmitoylation in the nucleus: a little fat around the edges. Nucleus 3, 251–255 (2012).

53. Park, S. et al. Palmitoylation controls the dynamics of budding-yeast heterochromatin via the telomere-binding protein Rif1. Proc Natl Acad Sci U S A 108, 14572–14577 (2011).

54. Fontana, G.A. et al. Rif1 S-acylation mediates DNA double-strand break repair at the inner nuclear membrane. Nat Commun 10, 2535 (2019).

55. Foti, R. et al. Nuclear Architecture Organized by Rif1 Underpins the Replication-Timing Program. Mol Cell 61, 260–273 (2016).

56. Xu, L. & Blackburn, E.H. Human Rif1 protein binds aberrant telomeres and aligns along anaphase midzone microtubules. J Cell Biol 167, 819–830 (2004).

57. Bhowmick, R. et al. The RIF1-PP1 Axis Controls Abscission Timing in Human Cells. Curr Biol 29, 1232–1242 e1235 (2019).

58. Germe, T., Miller, K. & Cooper, J.P. A non-canonical function of topoisomerase II in disentangling dysfunctional telomeres. EMBO J 28, 2803–2811 (2009).

59. Maciejowski, J., Li, Y., Bosco, N., Campbell, P.J. & de Lange, T. Chromothripsis and Kataegis Induced by Telomere Crisis. Cell 163, 1641–1654 (2015).

60. Nassour, J. et al. Autophagic cell death restricts chromosomal instability during replicative crisis. Nature 565, 659–663 (2019).

61. Masamsetti, V.P. et al. Replication stress induces mitotic death through parallel pathways regulated by WAPL and telomere deprotection. Nat Commun 10, 4224 (2019).

62. Tang, S., Stokasimov, E., Cui, Y. & Pellman, D. Breakage of cytoplasmic chromosomes by pathological DNA base excision repair. Nature 606, 930–936 (2022).

63. Bakhoum, S.F. et al. Chromosomal instability drives metastasis through a cytosolic DNA response. Nature 553, 467–472 (2018).

64. Nassour, J. et al. Telomere-to-mitochondria signalling by ZBP1 mediates replicative crisis. Nature (2023).

65. Liu, S. et al. Nuclear envelope assembly defects link mitotic errors to chromothripsis. Nature 561, 551–555 (2018).

66. Moreno, S., Klar, A. & Nurse, P. Molecular genetic analysis of fission yeast Schizosaccharomyces pombe. Methods Enzymol 194, 795–823 (1991).

67. Bahler, J. et al. Heterologous modules for efficient and versatile PCR-based gene targeting in Schizosaccharomyces pombe. Yeast 14, 943–951 (1998).

68. Roguev, A., Wiren, M., Weissman, J.S. & Krogan, N.J. High-throughput genetic interaction mapping in the fission yeast Schizosaccharomyces pombe. Nat Methods 4, 861–866 (2007).

69. Roguev, A., Xu, J. & Krogan, N. Genetic Interaction Mapping in Schizosaccharomyces pombe Using the Pombe Epistasis Mapper (PEM) System and a ROTOR HDA Colony Replicating Robot in a 1536 Array Format. Cold Spring Harb Protoc 2018 (2018).

70. Valente, L.P. et al. Myb-domain protein Teb1 controls histone levels and centromere assembly in fission yeast. EMBO J 32, 450–460 (2013).

